# An olfactory receptor gene underlies reproductive isolation in perfume-collecting orchid bees

**DOI:** 10.1101/537423

**Authors:** P. Brand, I. A. Hinojosa-Díaz, R. Ayala, M. Daigle, C. L. Yurrita Obiols, T. Eltz, S. R. Ramírez

## Abstract

Speciation is facilitated by the evolution of reproductive barriers that prevent or reduce hybridization among diverging lineages. However, the genetic mechanisms that control the evolution of reproductive barriers remain elusive, particularly in natural populations. We identify a gene associated with divergence in chemical courtship signaling in a pair of nascent orchid bee lineages. Male orchid bees collect perfume compounds from flowers and other sources to subsequently expose during courtship display, thereby conveying information on species identity. We show that these two lineages exhibit differentiated perfume blends and that this change is associated with the rapid evolution of a single odorant receptor gene. Our study suggests that reproductive isolation evolved through divergence of a major barrier gene involved in chemically mediated pre-mating isolation via genetic coupling.

---

Speciation, the formation of new species from a single ancestral species, is facilitated by the emergence of reproductive barriers between lineages and is considered the most fundamental process in the generation of biological diversity (*1, 2*). While a growing number of studies have revealed that recently formed species often exhibit marked divergence across multiple genomic regions (*3*), the role of these genomic ‘islands of divergence’ in reproductive isolation remains controversial (*4–7*). Even when specific genomic regions can be associated with reproductive isolation, they usually encompass hundreds of genes of unknown function. As a result, few studies have successfully linked specific genetic loci to reproductive barrier traits or determined how they contribute to the speciation process (*7*–*9*). Here we combine a large-scale population level approach with high-resolution genome-wide diversification analyses to identify the genetic basis of a phenotypic trait that likely controls reproductive isolation in a pair of orchid bee species.

Male orchid bees actively collect volatile chemical substances from floral and non-floral sources to concoct highly species-specific perfume blends (*10*–*12*), which they subsequently expose during ritualized courtship displays (Fig. 1) (*13, 14*). The exact type of information conveyed remains unknown, but perfumes are clearly involved in mating behavior and species recognition (*15*). Because orchid bees acquire volatile chemicals directly from the environment, their olfactory system is critical for both perfume concoction by males and perfume detection by females, which effectively creates a strong linkage between male trait and female preference. Thus, changes in genes underlying olfactory perception can simultaneously alter the male perfume signal and the female perfume preference through pleiotropic effects (*16*), a scenario that could lead to the rapid evolution of assortative mating (*17, 18*). We hypothesize that the differentiation of chemosensory genes drives rapid shifts in chemical perfume composition during the speciation process, facilitated by genetic coupling of male trait and the associated female preference in orchid bees.

**Fig. 1.**
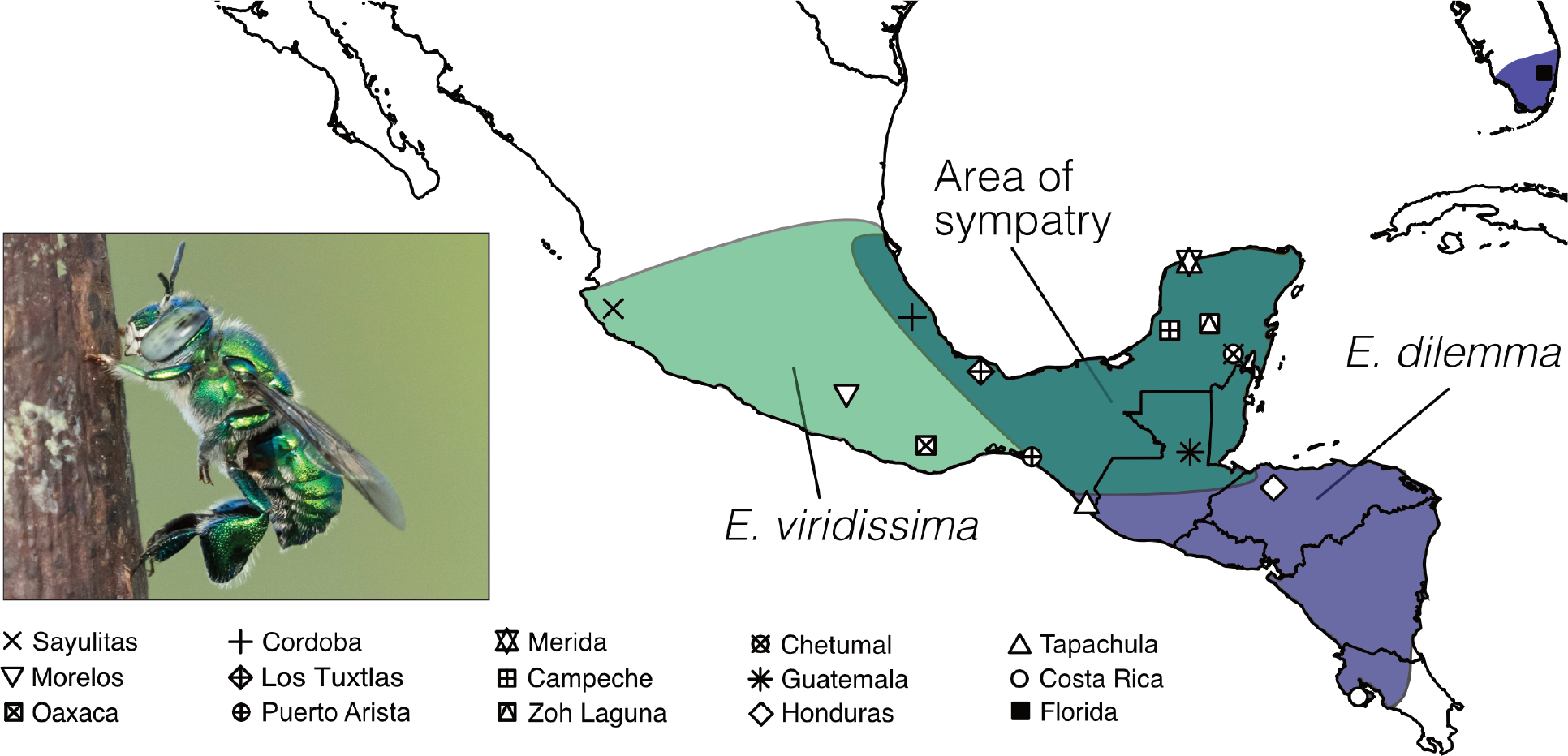
Distribution range of Euglossa dilemma and *E. viridissima*. Bees were collected in 15 sampling sites (Table S1) throughout the distribution ranges of each species including both allopatric (blue: *E. dilemma*, light green: *E. viridissima*) and sympatric populations (dark green). Photograph shows *E. dilemma* male perching during perfume display.

To test whether perfume composition evolves rapidly during species formation, we conducted a population-level analysis of perfume chemistry in a pair of orchid bee lineages (*Euglossa dilemma* and *E. viridissima*) that diverged ~150,000 years ago (*19*). We collected male bees across the entire geographical range of each lineage throughout Central America (Fig. 1, Table S1-S2) and analyzed the perfume chemistry of 384 individuals via gas chromatography-mass spectrometry (Table S2). A non-metric multidimensional scaling analysis revealed strong differentiation of perfumes into two distinct lineage-specific chemical phenotypes independent of geography (Fig. 2a, ANOSIM R=0.8, p=0.001). This pattern was driven by both quantitative and qualitative differences of perfume chemistry and held true when using either the entire set of compounds or the 40 most prevalent compounds (Fig. S1, Supplementary Text). This observation supports the hypothesis that those compounds collected by a high number of individuals play a critical role in perfume specificity and private signaling in orchid bees.

**Fig 2.**
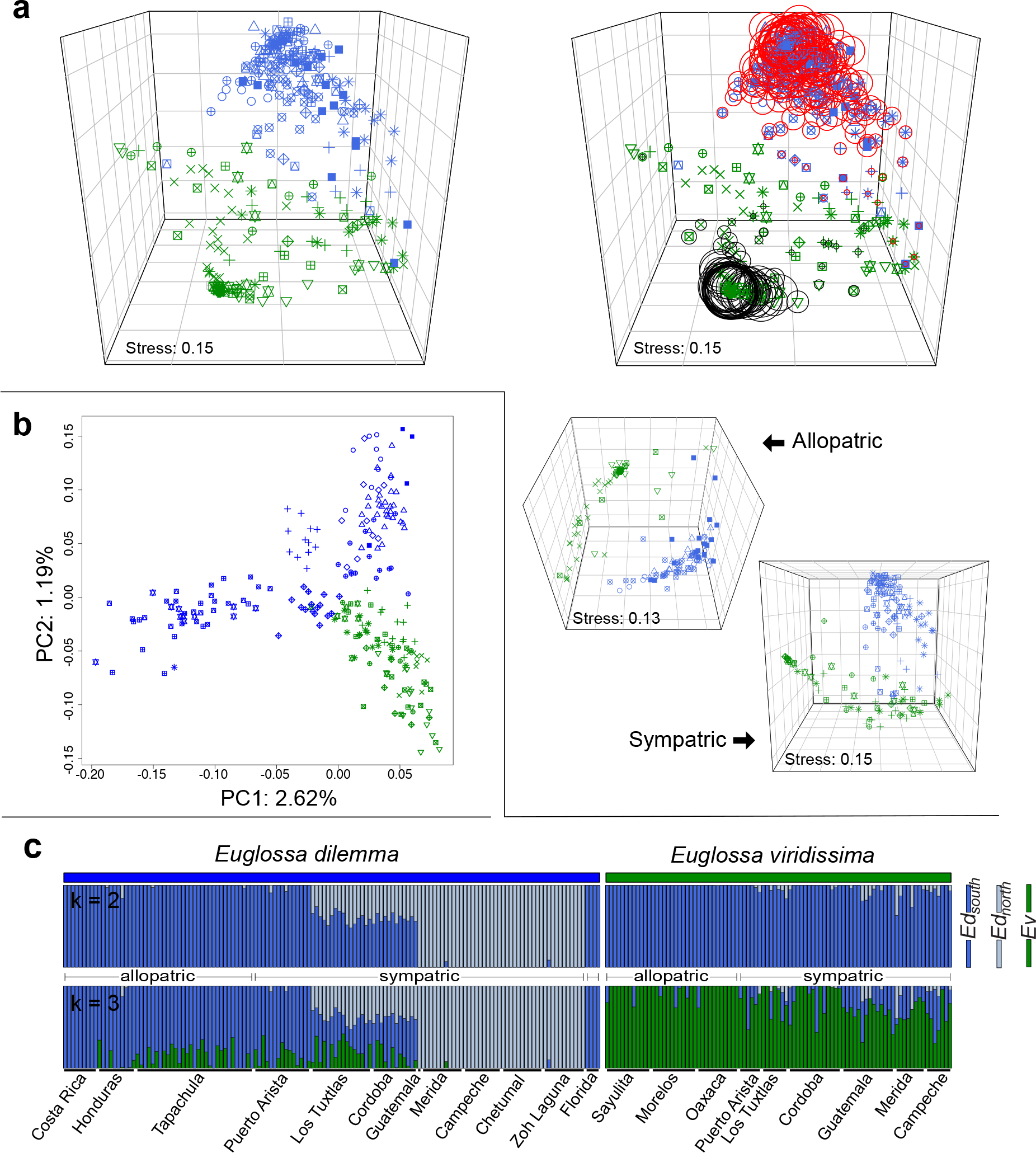
Phenotypic and genetic differentiation of *E. dilemma* and *E. viridissima*. (**a**) Perfume phenotypes were species-specific (upper left) independent of geography or co-occurrence (below) mainly due to the relative quantity of the major components HNDB (red circles), and L97 (black circles) as indicated by circle size (upper right). (**b**) *E. dilemma* (blue) and *E. viridissima* (green) are genetically differentiated over two PC axes explaining less than 4% of the genetic variation indicating that genetic differentiation is low. (**c**) Populations within *E. dilemma* (k=2) were separated before species (k=3) in a genetic clustering analysis, supporting population structure within *E. dilemma* (Fig. S6) and low interspecific genetic differentiation. Several individuals drew ancestry from multiple genetic lineages suggesting admixture.

We found that the most striking difference in perfume chemistry between *E. dilemma* and *E. viridissima* was the presence of two lineage-specific compounds that are highly prevalent and present in high relative abundance. The compound HNDB (2-hydroxy-6-nona-1,3-dienyl-benzaldehyde, (*16*)) was only present in perfume blends of *E. dilemma*, and the compound L97 (lactone-derivative of linoleic acid, (*20*)) was only present in perfume blends of *E. viridissima* (Fig. S2-S3, Table S3). These two molecules accounted for the highest average proportion of overall perfume content per species (relative abundance HNDB: 55%, L97: 37%, Table S4) and together contributed to 46.3% of the chemical differentiation between *E. dilemma* and *E. viridissima* (SIMPER analysis, Table S5).

Remarkably, perfume chemistry is the only trait that allows reliable identification and separation of these lineages. Close examination of morphological traits revealed that the number of teeth on the male mandible differs between the two lineages, with most but not all individuals segregating into two groups (*19*). Males of *E. dilemma* always have three mandibular teeth, whereas males of *E. viridissima* are polymorphic for the number of teeth with 89.4% of males exhibiting two teeth and the remaining fraction (10.6%) exhibiting three teeth that are hardly distinguishable from *E. dilemma* (Fig. S4, Supplementary Text). Together, these results demonstrate that species-specificity in perfume chemistry evolved rapidly through changes of few major compounds. These observations are consistent with the hypothesis that perfume chemistry is a mating recognition signal that functions as a pre-mating reproductive barrier among orchid bee lineages.

To contrast the divergence we observed in perfume chemistry with genetic differentiation between *E. dilemma* and *E. viridissima*, we genotyped 232 males sampled from across their geographic ranges (Fig. 1, Table S2). A principal components analysis of genetic variance (PCA) based on 16,369 single nucleotide polymorphisms (SNPs) revealed that these lineages are genetically distinct in both allopatric and sympatric populations (Fig. 2b, Fig. S5). *E. dilemma* and *E. viridissima* were not separated over a single PC axis (Fig. 2b, Fig. S6, Supplementary Text), which is consistent with a scenario of incomplete genome-wide separation between species. This observation was further supported by a genetic clustering analysis that first separated geographically distinct populations within *E. dilemma* before it separated species (ADMIXTURE, Fig. 2d, Fig. S7, Supplementary Text). In fact, this analysis revealed the existence of three main genetic lineages including 1) the entire *E. viridissima* population (*Ev*), 2) a southern *E. dilemma* population (*Ed*_*south*_), and 3) a northern *E. dilemma* population (*Ed*_*north*_). We also identified admixture between *E. dilemma* and *E. viridissima* (*f*_*4*_-test: 0.001, z: 2.7, p < 0.007, Table S6, Supplemental Text), as suggested by individuals that draw ancestry from multiple genetic lineages in the clustering analysis (Fig. 2d). These results show that while perfume chemistry evolved into two discrete chemical phenotypes, genetic differentiation is incomplete and low between *E. dilemma* and *E. viridissima* (pairwise *F*_ST_: 0.04 to 0.1, Table S7). These results support the hypothesis of ongoing gene flow during the early stages of speciation.

Next, to identify the genetic basis of perfume differentiation and reproductive isolation, we performed a genome-wide scan of divergence between *E. dilemma* and *E. viridissima* on the basis of 30 genomes from the three genetic lineages (N=10 for *Ed*_*north*_, *Ed*_*south*_, *Ev*, each, Fig. S8, Table S2, Supplemental Text). Genomic regions that contribute to a species-specific reproductive barrier trait should show high levels of differentiation due to diversifying selection between lineages but not within lineages. We capitalized on the fact that *E. dilemma* exhibits population structure to distinguish and identify regions of high differentiation between *E. dilemma* and *E. viridissima* but not within *E. dilemma*. Therefore, we estimated the net interspecific differentiation, which is calculated by subtracting the intraspecific *F*_ST_ from the interspecific *F*_ST_ (Δ*F*_ST_), for non-overlapping 50 kilobase (kb) windows across the genome (*21*). The resulting windows of elevated Δ*F*_ST_ (>99^th^ percentile) were clustered into eight distinct outlier peaks of varying size (0.05 to 1.7 Mb, Fig. 3b, Fig. S9, Table S8) that revealed elevated levels of genetic linkage compared to non-outlier windows (Mann-Whitney U Test, p<0.0001).

**Fig 3.**
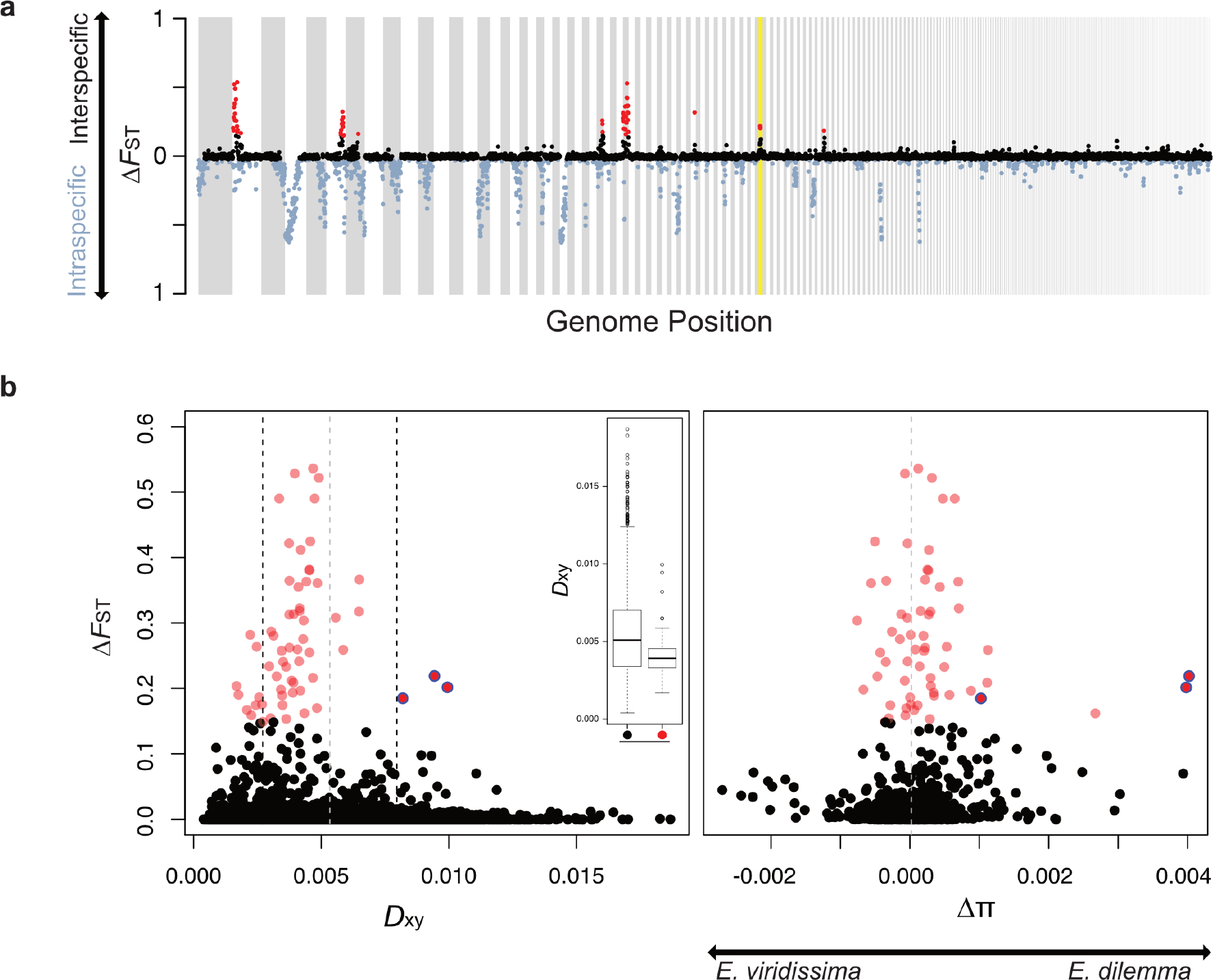
Whole-genome differentiation. (**a**) Eight regions of the genome revealed higher interspecific (black) than intraspecific (blue) differentiation (Δ*F*_ST_ >99^th^ percentile red). (**b**) Interspecific divergence (*D*_xy_) was negatively correlated with Δ*F*_ST_ (left panel, r=−0.06, p=0) and significantly reduced in outlier windows (red) in comparison to non-outliers (black, inlet, Mann-Whitney U test, p<0.0001). Only two of three Δ*F*_ST_ outlier windows (circled blue) that revealed increased *D*_xy_ also had and a net differential of intraspecific nucleotide diversity (Δπ) skewed towards *E. dilemma* (right panel), a pattern expected in genomic regions evolving under positive selection. Both correspond to the same outlier peak (yellow background in (**a**)). Grey dashed lines: mean *D*_xy_ and Δπ. Black dashed lines: one standard deviation of mean *D*_xy_.

While these genomic regions are likely to have evolved under linked selection, we found that absolute sequence divergence (*D*_xy_) was significantly reduced in Δ*F*_ST_ outlier windows (Mann-Whitney U Test, p<0.0001, Fig. 3c), suggesting that genetic differentiation in most outlier regions was not driven by diversifying selection and thus most likely unrelated to selection for species differences (*4, 5, 22*). Notwithstanding this general trend, we identified three outlier windows with elevated values of both *ΔF*_*ST*_ and *D*_*xy*_ (Fig. 3c), two of which exhibited a highly skewed differential in nucleotide diversity towards *E. dilemma* (Δπ, Fig. 3c), suggesting strong unilateral positive selection in this lineage. We identified signatures of an *E. dilemma*-specific selective sweep in one of these windows based on allele frequency spectra and haplotypes of the three distinct genetic lineages (Fig. 4a). We did not identify any additional species-specific sweeps (Fig. S10), highlighting that this 50 kb window contains a unique, locally restricted signature of positive selection in *E. dilemma*. This observation is congruent with strong and recent selective forces driving the divergence between *E. dilemma* and *E. viridissima* and suggests that the identified genomic region harbors loci mediating reproductive isolation between these nascent bee lineages.

**Fig 4.**
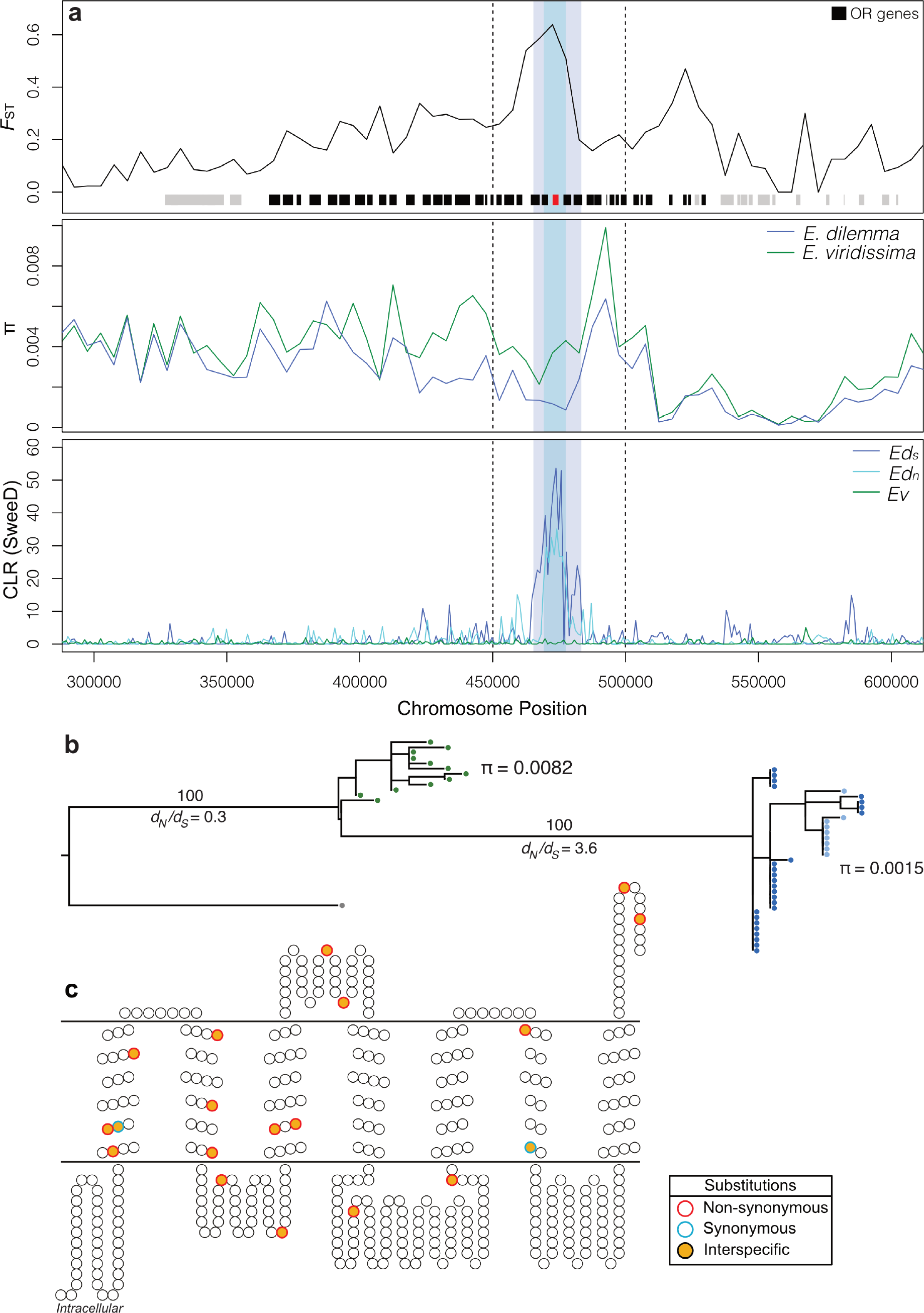
Odorant receptor (OR) gene *OR41* evolved through a species-specific selective sweep. (**a**) The only species-specific selective sweep identified was located within an *F*_ST_ outlier window (dashed lines) overlapping with a high interspecific difference in π in the middle of a tandem array containing 37 OR genes. High composite likelihood ratios (CLR, bottom) within *Ed*_*north*_ (light blue) and *Ed*_*south*_ (dark blue) but not in *Ev* (green) indicate a selective sweep shared by both *E. dilemma* lineages that overlap with *OR41* in the center (shaded regions). (**b**) A Maximum Likelihood phylogeny of *OR41* (N=47 individuals) demonstrates that genotypes are species-specific. π was five times lower in *E. dilemma* in comparison to *E. viridissima*. A *d_N_/d_S_* analysis of species-specific genotypes with five outgroup species (grey dot) indicates positive selection on the *E. dilemma* branch (*d_N_/d_S_*=3.6), but purifying selection of the ancestral genotype in *E. viridissima* (*d_N_/d_S_*=0.3). Bootstrap support for tested branches is indicated. (**c**) 17 of 19 substitutions mapped on the predicted membrane topology of the OR41 protein were non-synonymous.

Close inspection of the selective sweep region revealed the presence of 14 genes (Fig. 4a), all of which belong to the odorant receptor (OR) gene family and are located within a large ~170 kb tandem array of 39 ORs (*23*). The OR gene family is the largest chemosensory gene family in insects and is integral to the sensory detection of odorant compounds including pheromones (*24, 25*). Olfactory tuning is controlled by the OR protein sequence and therefore amino acid substitutions can shift odorant binding properties and sensory perception (*26, 27*). To identify the specific genetic targets of divergent selection, we mapped loci within the tandem array. We found that the region containing the selective sweep overlapped with both elevated interspecific *F*_ST_ values and reduced nucleotide diversity (π) in *E. dilemma* centered around a single OR gene, *OR41*, that we previously identified as divergent between *Ed*_north_ and *Ev* (*28*) (Fig. 4a). This suggests that *OR41* evolved under strong positive selection in the common ancestor of *E. dilemma* after or during the split between *E. dilemma* and *E. viridissima*. Re-sequencing of *OR41* confirmed these results (N=47, Fig. 4b, Table S9-S10, Supplementary Text), and revealed that the protein coding sequences were fixed for 19 substitutions between species, 17 of which were non-synonymous leading to changes in the amino acid sequence of the resulting protein (Fig. 4c). A comparison with distantly related *Euglossa* species demonstrated that all fixed substitutions were derived (Fig. S11) and evolved under strong positive selection in *E. dilemma* (*d_N_/d_S_* = 3.6, χ^2^ =16.1, p <0.0001, Table S11, Supplemental Text) but not *E. viridissima* (*d_N_/d_S_* = 0.3), suggesting that strong selective forces fixed amino acid substitutions and possibly drove odorant perception changes in *E. dilemma*. Future studies should focus on elucidating the binding properties of this receptor and each allelic variant.

Our results show that a simple major phenotypic difference in a reproductive barrier trait between two lineages in the early stages of speciation is maintained despite low genetic differentiation and ongoing gene flow. Only strong selection can counteract such equalizing mechanisms, highlighting the adaptive value of the species-specific major perfume compounds in *E. dilemma* and *E. viridissima*. While genome-wide analyses often lack resolution to identify the genes that control barrier traits (*3, 6*–*9, 29*), we were able to identify a single genetic locus of adaptive interspecific divergence, leading to a unique opportunity to understand the genomic landscape of speciation on a fine genetic scale in a non-model system. Our findings provide a link between a discrete shift in perfume composition with a single olfactory receptor gene that evolved under strong positive selection, linking a chemosensory barrier trait with an olfactory gene. Perfume composition in orchid bees is intricately connected to the sense of smell (*16, 30, 31*). In fact, *E. dilemma* and *E. viridissima* are known to differ in the behavioral preference towards and the sensory detection of HNDB (*16*) the major perfume compound in *E. dilemma*. This observation lends support to the hypothesis that the 17 non-synonymous substitutions present in *OR41* in the *E. dilemma* lineage underlie functional changes in sensory perception between species. Accordingly, the data presented here are consistent with the genetic coupling of a reproductive trait and trait preference (*17, 18, 32*) that evolved through rapid divergent selection in a single gene leading to speciation.

## Acknowledgements

We thank K.W. McCravy for sharing the Honduras bee samples; T. Pokorny, J.J.G. Quezada-Euan, R. Medina, M.A. Arteaga, and C. Pozo for help with field work; J. Fong for help with morphometry; the Ward lab for support with imaging; and M. Servedio and the Ramirez lab for helpful discussion.

## Author contributions

Conceptualization: P.B., T.E., S.R.R.; Fieldwork: P.B., S.R.R., I.H.-D., C.L.Y.O., R.A.; Data generation: P.B., M.D.; Project administration, Data curation, Formal analysis, Investigation, Visualization: P.B.; Funding acquisition: P.B., T.E., S.R.R.; Writing – original draft: P.B.; Writing – review & editing: P.B., T.E., S.R.R. with input from all authors; Supervision: S.R.R. work.

## Data and materials availability

Raw sequence data are available through NCBI (BioProject XXX), GCMS data are available through Dryad (xxx).

