## Supplemental Materials for "An olfactory receptor gene underlies reproductive isolation in perfume-collecting orchid bees"

Brand et al.

### Supplemental Material

### Materials and Methods

#### Sampling

We sampled males of the two orchid bee species *Euglossa dilemma* and *E. viridissima* between 2014 and 2016 throughout the entire distribution area of each lineage (Table S1) using chemical baits following Brand et al. (2015). Sampling and export of bees was performed with the necessary permits issued to Santiago Ramírez (Costa Rica, permit 050-2013-STNAC by the Ministerio del Ambiente y Energía), Ismael Hinojosa-Diaz (Mexico, permit SGPA/DGVS/09586/15 by the Secretaría de Medio Ambiente y Recursos Naturales), and Carmen Lucía Yurrita Obiols (Guatemala, permit 2756/2016 by the Consejo Nacional de Areas Protegidas). We dissected males in the field directly following collection and stored both hindlegs in 500 µL hexane for perfume extraction, while the rest of the body was preserved in 95% ethanol for subsequent morphological and genetic analyses. Perfume extracts and body tissue were stored at -20° C until analyzed.

#### Perfume analysis

*Chemical analysis*

We analyzed the perfume composition of 384 individuals (Table S2) using gas chromatography- mass spectrometry (GC-MS) with an Agilent 7890B GC fitted with a 30 m X 0.25 mm x 0.25 µm HP-5 Ultra Inert column coupled with an Agilent 5977A MS (Agilent Technologies) housed at the University of California, Davis. Using an autosampler, we injected 1µL perfume extract splitless into the GC. Oven temperature was held at 60° C for 3 minutes and then increased by 3° C per minute until it reached 300° C. Finally, the oven temperature was kept at 315° C for 1 minute. Both injector and transfer line temperatures were kept at 250° C. Helium was used as carrier gas with a flow rate of 1.2 mL per minute. Electron impact mass spectra were obtained by scanning between 30 and 550 mass-to-charge ratio (m/z). GC-MS data were processed using the MassHunter GC/MS Acquisition software vB.07.00 (Agilent) and analyzed in OpenChrom v1.1.0 (Lablicate).

*Compound characterization*

We created a manual mass spectral database from chromatogram peaks in OpenChrom, which we used to cross-reference the chromatograms of all analyzed individuals. In order to match peaks from different chromatograms, a minimum of 95% overlap of mass spectra with the manual database was required. We updated the database recursively as new compounds were detected. Individual compounds in the manual database were characterized by comparing mass spectra against the NIST05 database using the NIST MS Search software v2.0 as well as other published mass spectra (Ramírez *et al.* 2010; Eltz *et al.* 2011; Pokorny *et al.* 2013). Chromatogram peaks were detected in OpenChrom using the first derivative peak detector with minimum signal-to-noise ratio set to 5 and a moving average window size set to 17. To determine total ion abundances of each peak, we integrated peaks in OpenChrom using the standard integrator. These steps were automatized to analyze all 385 samples in batch mode. Only peaks with an area ≥1% of the larges peak were included in downstream analyses. Peaks that corresponded to chemicals of endogenous origin such as cuticular hydrocarbons and other glandular secretions were identified via comparison to chromatograms from the labial glands, hindlegs of males hatched in captivity that had not collected perfume compounds, and previously generated species-specific mass spectral databases (Eltz *et al.* 2008; Ramírez *et al.* 2010); these compounds were removed from all subsequent analyses.

*Statistical analysis*

We manually aligned the perfume profiles of all 384 individuals (Table S2) using a combination of retention time and compound identity following Eltz *et al.* (1999). The resulting matrix containing absolute quantities (total ion currents) for each compound was used for all downstream statistical analyses. Individuals with less than 10 collected compounds and thus in the early stages of perfume collection were discarded from subsequent analyses leading to a final set of 306 individuals in the perfume dataset. Similarly, compounds identified in less than three individuals were removed from the analysis since these compounds likely represent molecules collected accidentally and are not likely involved in chemical signaling. We then transformed the absolute quantities of compounds to relative amounts per individual and analyzed the final perfume matrix in R following Ramirez et al. (2010). Briefly, we compared individual chemical profiles of the entire dataset as well as allopatric and sympatric subsets using three-dimensional non-metric multidimensional scaling (nMDS) analyses. Therefore we calculated a triangular distance matrix between individuals using the Bray-Curtis (BC) index of dissimilarity, which is insensitive to compounds absent in sample pairs. Based on this BC matrix, we computed 2- and 3-dimensonal nMDS plots with 50 iterations per run using the ecodist package v2.0.1 (Goslee and Urban 2007). Each analysis was run 10 times and convergence between runs was visually inspected. Only the 3D analysis was retained due to high stress values of the 2D nMDS analyses. To statistically assess whether perfume profiles are more dissimilar between than within species, we conducted an Analysis of Similarity (ANOSIM) test implemented in the vegan package v2.5.2 (Oksanen *et al.* 2018). We further estimated the relative contribution of each individual compound to the observed ordinal dissimilarities using the Similarity Percentage (SIMPER) method as implemented in vegan. Additional descriptive statistics and plots were produced in R.

**Mandible morphometrics**

We determined the number of mandibular teeth in 414 males collected throughout the distribution range of *E. dilemma* and *E. viridissima* (Table S2). A total of 130 individuals had two teeth, while the remaining 283 had three teeth on the mandible. To perform geometric morphometric analyses between *E. dilemma* and *E. viridissima*, we dissected the mandibles of 175 tridentate individuals (Table S2) from the head capsule and mounted them on a paper point using clear nail polish. With few exceptions, the left mandible was used. In few cases the left mandible was too damaged to mount, and therefore we used the right mandible (as indicated in Table S2) and subsequently digitally created a mirror image before it was included in the dataset. Mandibles were imaged using a Leica MZ 16A stereomicroscope with 100x magnification. We captured stacked photographs with a JVC KY-F57U camera mounted on the stereomicroscope and merged them using Auto-Montage Pro (Synoptics Ltd., Cambridge, England). We converted the resulting images to the thine-plate spline (TPS) format in tpsUtil (Rohlf 2004), which we subsequently used to set five landmarks corresponding to the three tips of each tooth and the two indentations in-between teeth (Figure S1) in tpsDig (Rohlf 2009). The resulting landmark data was analyzed with geomorph 3.0.7 (Adams and Otarola-Castillo 2013) in R by overlaying the landmarks of all individuals to identify species-specific geometric morphometric differences. A PCA of landmark shape variation was conducted using the plotTangentSpace function in geomorph.

**Population genetics**

<u>*DNA extraction and sequencing*</u>

To extract DNA, we dissected flight musculature from the thorax of individuals stored in 95% ethanol using sterile forceps, dried it for 1-2 hours at room temperature, deep froze it in liquid nitrogen, and then ground it on a tissue lyzer (Qiagen). DNA was then extracted using the Blood and Tissue Kit following the manufacturers instructions after an overnight proteinase K digestion step (Qiagen). The extracted DNA was used for genotype-by-sequencing (GBS) following Elshire *et al.* (2011). Briefly, the DNA of each individual was digested using the EcoT22I restriction enzyme, followed by barcode ligation. 95 individually barcoded samples were then pooled and PCR amplified, and finally size selected for ~300bp using AMPure bead cleanup. Libraries were then validated on a Bioanalyzer high-sensitivity DNA chip (BioRad) and sequenced on a HiSeq2100 (Illumina) in 150 bp single-end mode. Three individuals were run on each of three lanes to correct for potential batch effects.

<u>*Statistical analyses*</u>

Sequencing reads were de-multiplexed using Stacks v1.47 (Catchen *et al.* 2013) and mapped to the *E. dilemma* reference genome (Brand *et al.* 2017) using Bowtie 2 (Langmead and Salzberg 2012). Since male Hymenopterans are haploid, we discarded all reads that mapped to more than one region of the genome to exclude repetitive loci. Nucleotides were called when a locus was sequenced in at least 50% of all individuals to a minimum read depth of four reads. SNPs were pruned if they were in linkage disequilibrium of $r^2 \geq 0.2$ or had more than two alleles using plink v1.9 (Purcell *et al.* 2007). After initial filtering, we excluded all individuals with less than 5000 called SNPs and repeated the filtering steps above on the final set of 232 individuals. The resulting pre-processed SNP set was then used for all downstream analyses.

To visualize genetic structure among populations and species we performed a Principal Components Analysis of genetic variance (PCA; Price *et al.* 2006) in the R package SNPRelate (Zheng *et al.* 2012). Independent PCAs were performed on the entire set of individuals as well as on subsets of all sympatric sampling sites. In addition, we performed ancestry estimation in ADMIXTURE (Alexander *et al.* 2009). For both analyses the SNP set was pruned to a minimum allele frequency (MAF) of 5%. To estimate admixture proportions for each individual in the dataset, we used ADMIXTURE based on SNPs called in at least 75% of individuals. We estimated levels of genetic structure for $k = 1$ to 10 jointly for all individuals. Each run was repeated 10 times with random seeds including cross-validation (CV) error estimation for each level of *k*. We inferred the estimated number of genetic clusters using the mean CV error.

We inferred the fixation index ($F_{ST}$) between genetic clusters in SNPRelate.

In order to test for 'treeness' of the phylogeny including sympatric and allopatric populations of *E. dilemma* and *E. viridissima* we used the $f_4$-test (Keinan *et al.* 2007) as provided as part of the treemix software package v1.12 (fourpop; Pickrell and Pritchard 2012). The $f_4$-test is based on the null hypothesis that the evolutionary history of four pre-defined lineages can be explained by a simple bifurcating phylogeny. In case the null hypothesis is rejected, it suggests that other mechanisms such as admixture or incomplete lineage sorting of ancestral population structure were involved in the evolution of the tested lineages. We performed the $f_4$-test to test for the potential of gene flow between *E. dilemma* and *E. viridissima* in sympatric areas by running $f_4$(Edil_allo, Edil_sym; Evir_allo, Evir_sym). We excluded the individuals from Florida and Puerto Arista for this analysis, because they were unusual. The Florida population has lower nucleotide diversity than all other sampling sites (data not shown) likely due to a recent bottleneck after its introduction (Zimmermann *et al.* 2011) and Puerto Arista individuals are sympatric but genetically belong to the otherwise allopatric $Ed_{south}$ lineage (see main text and below) and thus could not unequivocally assigned to either the allopatric or sympatric group. The SNP set used was pruned to unlinked SNPs called in at least 75% of individuals with a MAF of 0.05 (1,399 SNPs total). Standard errors were estimated using each SNP during jackknifing in the fourpop program (-k option set to 1).

**Population genomics**

<u>*Sequencing*</u>

To conduct genome-wide analyses of population and species differentiation, we sequenced the whole genomes of 10 individuals of each of the three genetic lineages identified ($Ed_{north}$, $Ed_{south}$, *Ev*). DNA from individuals with known GBS genotype was used for whole-genome library preparation using an adapted diluted Nextera DNA Sample Preparation procedure (New England Biolabs) following Baym *et al.* (2015) to a minimum genome-wide read-depth of 5x.

<u>*Statistical analysis*</u>

Reads were mapped to the *E. dilemma* reference genome (Brand *et al.* 2017) using bwa-mem (Li 2013) and SNPs were called with GATK v4.0.1.2 (McKenna *et al.* 2010). Genome differentiation was analyzed in R using PopGenome (Pfeifer *et al.* 2014), due to its support for haploid datasets. We used a non-overlapping 50kb sliding window approach to estimate pairwise relative genetic differentiation ($F_{ST}$), absolute sequence divergence ($D_{xy}$), and nucleotide diversity (π) for each window between all three genetic lineages. In addition, we calculated linkage disequilibrium (LD, $r^2$) within the same 50kb windows using vcftools. Additional descriptive statistics were produced using plink and base R.

In order to filter genomic windows of elevated interspecific divergence, we calculated the net interspecific differentiation ($\Delta F_{ST}$, Vijay *et al.* 2016) by subtracting intraspecific diversity within *E. dilemma* from interspecific diversity between *E. dilemma* and *E. viridissima* ($\Delta F_{\mathrm{ST}} = F_{\mathrm{ST}}$ [*Ev* vs. *Ed*] - $F_{\mathrm{ST}}$ [$Ed_{north}$ vs. $Ed_{ssouth}$]) since genetic regions that are involved in species delimitation are expected to be differentiated between but not within species. We then filtered the >99$^{\mathrm{th}}$ percentile $\Delta F_{\mathrm{ST}}$ regions as outliers of interspecific differentiation. In order to identify windows with $\pi$ values biased towards one species, we contrasted $\pi$ between species by subtracting $\pi_{Ed}$ from $\pi_{Ev}$ to calculate the net differential in intraspecific nucleotide diversity ($\Delta\pi$) between the two species. If $\Delta\pi$ equals 0, $\pi$ is indifferent between species in the corresponding genomic window. Correlations of the different statistics and outlier tests were performed in R.

An increase in interspecific differentiation ($\Delta F_{\mathrm{ST}}$) in combination with a reduction in nucleotide diversity in one species (leading to a highly skewed $\Delta\pi$) is indicative of a selective sweep. Additionally, to ascertain candidate regions, we performed two independent tests for selective sweep signatures in all $\Delta F_{\mathrm{ST}}$ outlier windows using 1) SweeD (Pavlidis *et al.* 2013) and 2) hapFLK (Fariello *et al.* 2013). These two methods identify selective sweeps based on two different types of data including either the allele frequency spectra (SweeD) or haplotype information (hapFLK) of genomic regions. Due to their different algorithms, the two methods have different sensibilities in the detection of selective sweeps ranging from the early onset of selective pressures (hapFLK) to post-fixation of the beneficial allele (SweeD) with overlap in their ranges of detectability in-between (Vitti *et al.* 2013; Weigand and Leese 2018). We ran hapFLK with k=5 clusters and the default of 20 model fits (--nfit) on all scaffolds carrying $\Delta F_{\mathrm{ST}}$ outlier windows. The optimal number of clusters was calculated following the fastPHASE cross-validation method (Scheet and Stephens 2006) as implemented in the imputeqc R package (Khvorykh 2018). SweeD was run on the folded site frequency spectra using a sliding window size of 1000 bp. The $\Delta F_{\mathrm{ST}}$ outlier window that revealed a species-specific sweep pattern was then analyzed with a 10 kb non-overlapping sliding-window in SNPrelate as described above.

#### ***OR41* evolutionary history**

To infer the evolutionary history of *OR41* alleles across *E. dilemma* and *E. viridissima*, we re-sequenced the gene from individuals collected across the entire distribution ranges of both species using tiled Sanger sequencing. The *OR41* gene consists of 6 exons and 5 introns spanning a total of 2255 base pairs (Brand and Ramírez 2017). We designed a set of 4 tiled PCR primer-pairs spanning all exons and introns (Table S9) to amplify the whole gene in a total of 47 individuals (12 *Ev*, 9 $Ed_{north}$, 26 $Ed_{south}$; Table S10) using the following PCR program: initial 2:30 min at 94C followed by 28 cycles of 0:40 min at 94C, 0:40 min at 54C, and 0:50 min at 72C followed by a final 8:00 min at 72C. Individual fragments were aligned to the OR41 gene model derived from the *E. dilemma* reference genome (Brand and Ramírez 2017; Brand *et al.* 2017) and the open reading frames of either species (Brand *et al.* 2015) using mafft v7.215 (Katoh and Standley 2013) in the l-INS-I mode. Based on these alignments the OR41 gene sequence of each individual was reconstructed in Geneious v8 (Biomatters Ltd.).

After the reconstruction of *OR41* genotypes, we produced a multi-sequence alignment including all individuals in mafft and analyzed it in MEGA v5 (Tamura *et al.* 2013). We estimated π within each species for the entire gene. Subsequently, substitutions in the open reading frame were identified visually and defined as fixed between species when all individuals of one species had a nucleotide different from all individuals of the other species. Otherwise, a substitution was defined polymorphic. Similarly, we visually identified substitutions as non-synonymous or synonymous based on the open reading frame. We then mapped the substitutions to the predicted membrane topology (Brand *et al.* 2015).

To reconstruct the *OR41* evolutionary history, we inferred a phylogenetic tree based on the entire ORFs of the 47 sequenced individuals and the publicly available ORFs of *E. imperialis, E. flammea,* and the more distantly related orchid bee *Eufriesea mexicana* (Brand and Ramírez 2017) as outgroup. For tree inference, we estimated a maximum likelihood tree in RaxML following Brand and Ramirez (2017), including 1000 bootstraps.

To test for selection along the branches leading to the respective species, we produced a gene phylogeny based on the consensus sequence for each of the two species together with *E. imperialis, E. flammea,* and *Ef. mexicana* as outgroup. The tree was used for a $d_N/d_S$ test using codeml in PAML v4.6 (Yang 2007). Therefore, we estimated the likelihood of a model allowing for two or more $d_N/d_S$ values over branches in the tree with *E. dilemma* as foreground branch (M1), and a null model allowing only a single $d_N/d_S$ for all branches (M0). We conducted a likelihood ratio test to test whether the model with branch variation is more likely than the null model ($\Delta = 2(\ln(M1)-\ln(M0))$ with Δ approximating a $\chi^2$ distribution with one degree of freedom). Only fixed differences between *E. dilemma* and *E. viridissima* were taken into account to prevent overestimation of $d_N/d_S$ values (Kryazhimskiy and Plotkin 2008; Mugal *et al.* 2014).

### Supplementary Text

#### Perfume analysis

Perfume complexity

Individual perfume blends contained between 0 and 95 compounds with a mean of 29 ± 21 compounds (*E. dilemma*: 33 ± 22, *E. viridissima*: 24 ± 17). Of the 384 individuals analyzed, 306 had collected at least 10 compounds and were retained for all downstream analyses. In this filtered dataset the average per-capita number of compounds was 35 ± 19 (*E. dilemma*: 39 ± 20, *E. viridissima*: 29 ± 15). This is similar to previous analyses in *E. dilemma* (Ramírez *et al.* 2010). In total, we identified 391 different compounds of exogenous origin. Of these, 308 compounds (79%) were identified in less than 10% of all individuals. This set of low-prevalence compounds (present in <10% of individuals) was highly similar when each species was analyzed separately. Overlap of low-prevalence compounds between the total dataset and *E. dilemma* and *E. viridissima* subsets was 99% and 92%, respectively. The few compounds that were not overlapping and thus present in >10% of individuals in one or both subsets were present in no more than 17% of individuals in either species. This indicates that the set of low-prevalence compounds identified based on the entire dataset is highly representative of low-prevalence compounds in either species. It is unlikely that these low-prevalence compounds contribute to the signaling character of the perfume blend. To the contrary, it is more likely that low-prevalence compounds represent non-target compounds incorporated into perfumes by accident or because they co-occur at sources with core perfume compounds. This makes sense, because male orchid bees use a lipoid secretion from the labial gland to dissolve and collect all target compounds independent of the compounds mass or chemical class (Eltz *et al.* 2007). Since these secretions can act as solvents of a wide range of nonpolar chemicals, it is highly likely that off-target chemicals get incorporated as well during scent collection, leading to chemical background noise. Remarkably, even though male orchid bees obtain perfume compounds from flowers, fungi, and other sources (Dressler 1982; Whitten *et al.* 1993), both lineages exhibited strong perfume-specificity across their geographic range, which include highly heterogeneous habitats ranging from tropical cloud forests in Cordoba to dry forests in Costa Rica. The high levels of similarity in low-prevalence compounds collected by *E. dilemma* and *E. viridissima* might indicate that males from both lineages collect their shared compounds from similar sources. However, it is also possible that non-target compounds are similar among different collection sources.

Species-specific compounds

We hypothesized that perfume compounds that play an important role in species-specific signaling would be completely absent or of low prevalence in one lineage but present in the perfumes of most individuals of the other lineage. To identify species-specific compounds with putative signaling function, we extracted the low-prevalence compounds identified in only one of the two species and contrasted those with the prevalence in the other species. Overall, we identified four compounds present in <10% of *E. dilemma* individuals but in ≥10% *E. viridissima* individuals and 27 compounds in the reciprocal comparison (Figure S2). Of these, only five were present in more than 50% of individuals in one species and <10% in the other. These included L97, which is present in the perfumes of only 3% *E. dilemma* males but in 61% *E. viridissima* males as well as four stereoisomers of HNDB present in 86% to 98% of all *E. dilemma* individuals but only in 0% to 2% *E. viridissima* males (Table S3). This result extends previous analyses based on the sympatric Yúcatan populations alone (Eltz *et al.* 2008; 2011; Pokorny *et al.* 2013) to the entire distribution range of both species, and supports the hypothesis that L97 and HNDB are the main species-specific perfume compounds.

Perfume blends are species-specific due to HNDB and L97

We performed nMDS analyses on all compounds as well as the subset of the 40 compounds found in the highest number of individuals in the dataset (high-prevalence compounds, Table S3). Our analysis revealed strong differentiation between *E. dilemma* and *E. viridissima* for both subsets (Figure S3, ANOSIM R = 0.8, p=0.001). This suggests that the chemical disparity between species is mostly due to compounds collected by most individuals of either species, further supporting the hypothesis that the majority of compounds identified represent chemical background noise introduced during environmental scent collection. This makes sense given the expectation that compounds with a function in perfume signaling are likely enriched in comparison to background noise in a cross-population sample like this.

The SIMPER analysis revealed that only 16 compounds contributed more than 1% to the chemical disparity between *E. dilemma* and *E. viridissima* (Table S5). HNDB explained the most with 24.3%, followed by L97 with 18.6%. Only two more chemicals explained >3% chemical disparity including eugenol with 5.5% and cis-β-ocimene with 5.4%. All 16 compounds contributing more than 1% to the chemical disparity are also among the 40 most prevalent compounds (Table S3) as well as the compounds with the highest relative abundance of total perfume content (Table S4) in the dataset. In terms of relative abundance, HNDB and L97 also are extreme outliers. A mean of 55% total perfume content in *E. dilemma* corresponds to HNDB and a total of 37% in *E. viridissima* corresponds to L97 (Table S4). Accordingly, HNDB and L97 are more than nine and four times more highly concentrated, respectively, than the next most relative abundant compound (cis-beta-ocimene for both lineages, Table S4). With the exception of these major compounds, all other compounds with >1% relative abundance revealed similar mean relative abundances in both lineages, but benzyl benzoate revealed a relative abundance of 6% in *E. dilemma* and 0.1% in *E. viridissima* (Table S4). Although we identified a few individuals with the major component of the other lineage in the perfume (four *E. dilemma* males with L97 and three *E. viridissima* males with HNDB), the maximum relative abundance of a ‘mismatched’ compound identified was 4.7% for L97 in the perfume of an *E. dilemma* male that also contained HNDB with a relative abundance of 59.8%. This suggests that the major compound of the other lineage might be accidentally collected on rare occasions and likely represents a non-target compound similar to the low-abundance compounds identified. Overall, these results indicate that HNDB and L97 are distinctively species-specific and each likely only actively collected by one of the two bee lineages.

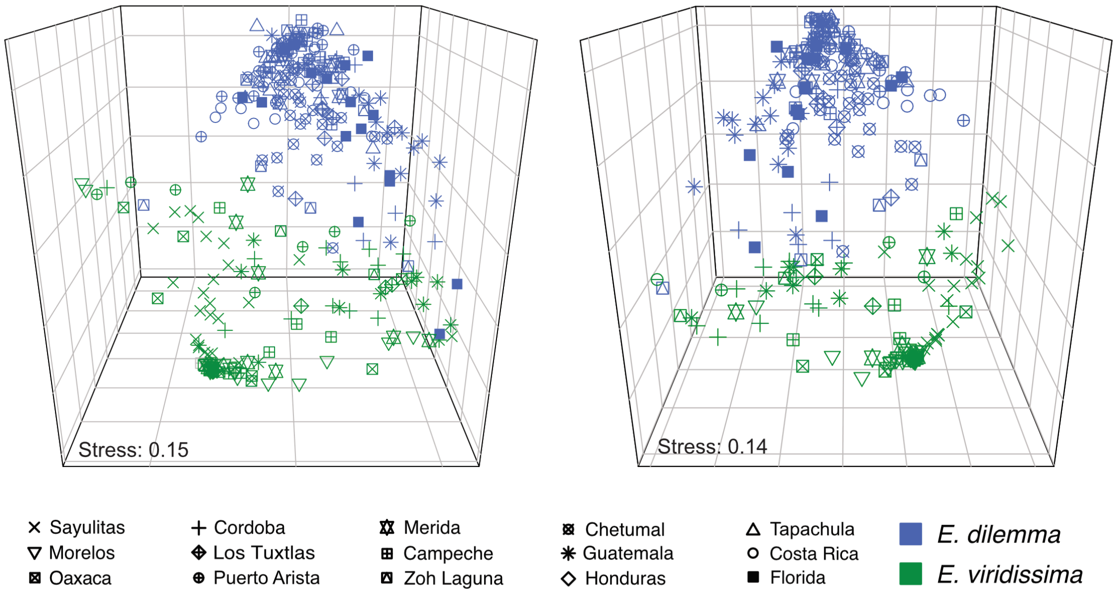

**Figure S1 Perfume differentiation of *E. dilemma* (green) and *E. viridissima* (blue).** Perfume phenotypes clustered species in nMDS analyses based on all perfume compounds (left) and the 30 most abundant compounds (right).

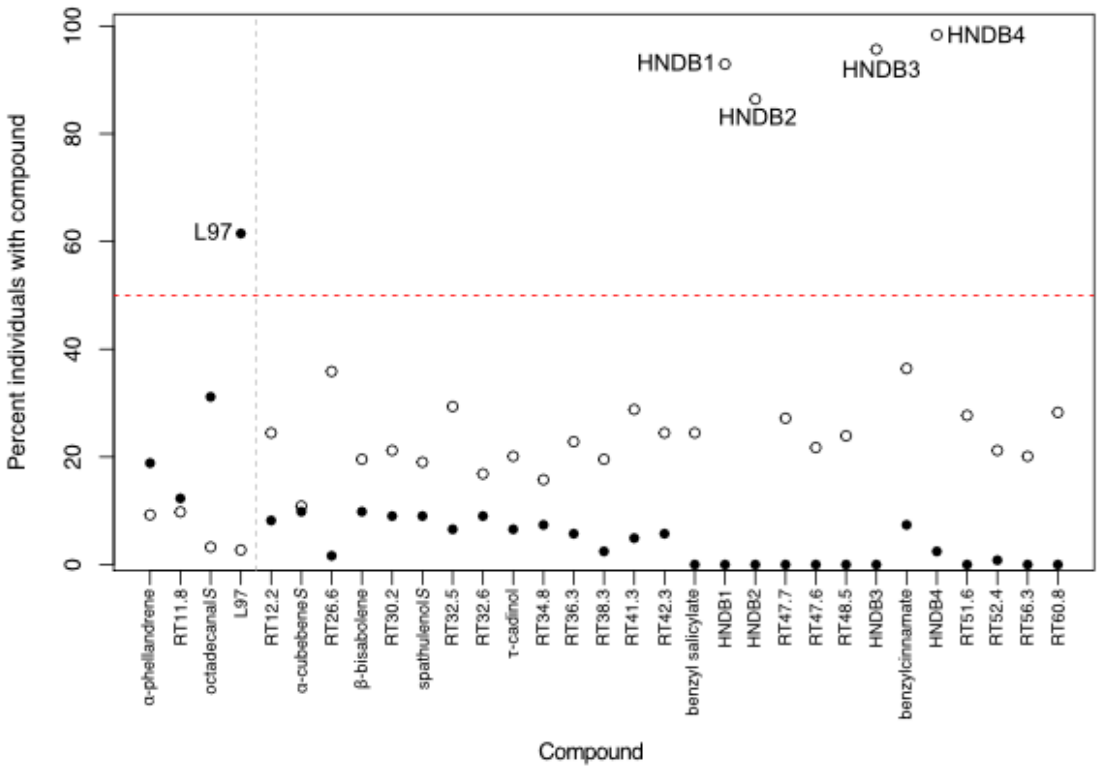

**Figure S2 Compounds of species-specific high abundance in *E. dilemma* (white) and *E. viridissima* (black).** Compounds present in perfumes of >10% of analyzed *E. viridissima* and <10% of analyzed *E. dilemma* individuals (left of grey dotted line) and the reciprocal comparison (right of the grey dotted line) revealed five compounds present in >50% of individuals of one lineage (above red dotted line) and <10% in the other. These are L97 of high abundance in *E. viridissima* and the four HNDB stereoisomers of high abundance in *E. dilemma*. An *S* appended to a compound name indicates compounds similar to the indicated.

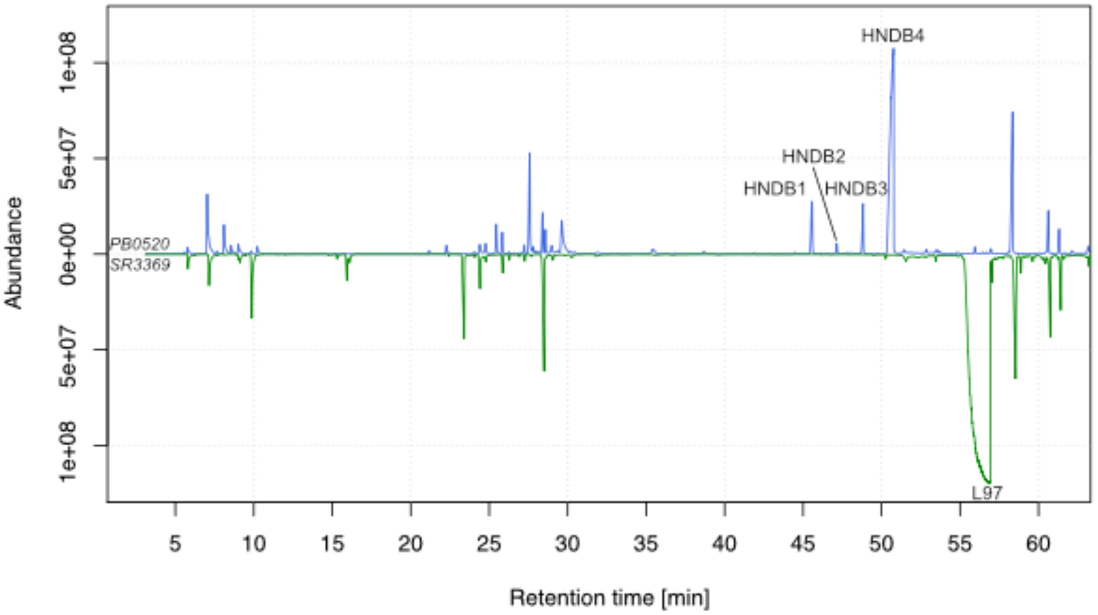

**Figure S3 Example chromatograms for *E. dilemma* (blue) and *E. viridissima* (green) perfumes highlighting the major compounds.** Peaks in the chromatogram represent single compounds present in the perfume of one individual of each lineage (PB0520 and SR3369). The higher a peak and the larger the area under its curve in relation to all other peaks the higher its relative abundance.

### Mandible morphometrics

Mandible dentation is the only morphological character identify so far that is differentiated between *E. dilemma* and *E. viridissima* with the potential for reliable determination in the field (Eltz *et al.* 2011). *E. dilemma* males have three mandibular teeth while most *E. viridissima* males have two teeth and occasionally three teeth, especially when sympatric with *E. dilemma*. Eltz et al. (2011) described mandible morphology of tridentate *E. viridissima* males as different from *E. dilemma* males based on the position of the central mandibular tooth, which they found to be shifted towards the basal tooth. However, their results were exclusively based on individuals collected in the northern part of the Yucatán peninsula, which only represents a small portion of the sympatric distribution. In order to test if these findings are applicable throughout the distribution ranges of the two lineages, we analyzed the mandibles of males we collected in 15 different sampling sites (Fig. 1, Table S1, Table S2).

Consistent with previous results (Eltz *et al.* 2011), all but three *E. viridissima* males collected from allopatric populations had two mandibular teeth (bidentate n=93, tridentate n=3), while all *E. dilemma* individuals had three teeth independent of co-occurrence with *E. viridissima* (n=287). In addition, a fraction of *E. viridissima* individuals in sympatry had three instead of the usual two teeth (bidentate n=34, tridentate n=12). All three allopatric tridentate *E. viridissima* individuals were collected in Oaxaca, which is close to the allopatry-sympatry boundary of the species (Fig. 1, main text). Our data suggest that tridentate *E. dilemma* males are more highly abundant in sympatry (26% of all individuals) than allopatry (3% of all individuals). However, the overall number of individuals collected in sympatry is comparatively small and thus estimates might change with the collection of a higher number of individuals. Our geometric morphometric analysis of 175 tridentate individuals of both lineages including 11 of the 12 tridentate *E. viridissima* individuals from the sympatric distribution range revealed that the mean position of the central mandibular tooth is shifted towards the basal tooth in *E. viridissima* tridentate males but that there is considerable overlap in tooth positions among individuals between the two lineages (Figure S1). This includes the sympatric area analyzed by Eltz et al. (2011).

Our results suggest that tridentate *E. viridissima* cannot be unequivocally distinguished from *E. dilemma* individuals in the field. However, the results indicate that individuals with a central mandibular tooth shifted towards the basal tooth are more likely to belong to *E. viridissima*. It is possible that tooth morphology is a labile trait varying among generations. The samples analyzed in Eltz et al. (2011) were all sampled before 2009, some of them more than 40 years ago and it might be possible that tooth morphology changed over short time scales. Further, it is possible that tridentate *E. viridissima* individuals are a result of hybridization between the two lineages. The fact that all but three tridentate *E. viridissima* individuals were found in sympatric but not allopatric populations lends support to this hypothesis. While this hypothesis remains to be tested, it would explain why the mandibular morphology of some tridentate *E. viridissima* is more similar to *E. dilemma* than others.

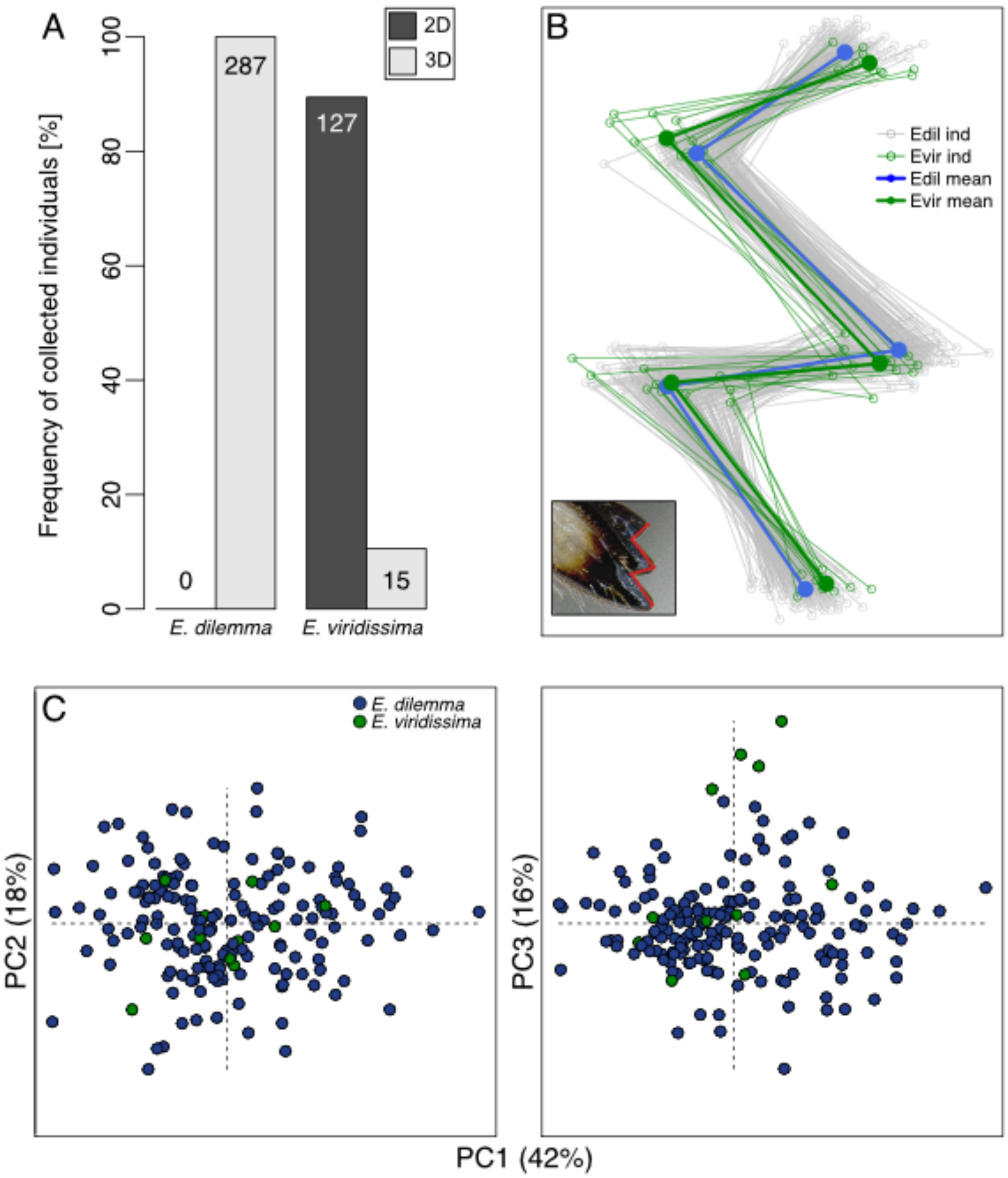

**Figure S1 Geometric morphological analysis of tridentate mandibles. A)** While all *E. dilemma* males collected were tridentate (3D), *E. viridissima* was polymorphic for the number of mandibular teeth with 10.6% of the analyzed males revealing tridentate and 89.4% bidentate (2D) mandibles, all of which were collected in sympatric populations. **B)** Individual landmarks of 175 tridentate mandibles were aligned for *E. dilemma* (N = 164, grey) and *E. viridissima* (N = 11, green) individuals (above left). Mean landmarks are indicated in thick lines and solid points for *E. dilemma* (green) and *E. viridissima* (blue) and suggest that overall the middle tooth in *E. viridissima* is shifted towards the basal tooth. The inlet indicates how landmarks were set on stacked photos of mandibles. **C)** PCA of mandible shape reveals no species-specific clustering over the first two PC axes (left). The third PC axis reveals clustering of four of the 11 *E. viridissima* individuals. Variance explained by the respective axes is indicated in brackets.

### Population genetics

Our GBS approach resulted in 16,369 SNPs sequenced in at least 50% of 232 males collected throughout the entire geographic ranges of *E. dilemma* and *E. viridissima* (Table S1, Table S2). Of these, 5,428 had a minor allele frequency (MAF) ≥ 0.05 indicating that most identified variants are rare.

*E. dilemma* population structure reveals the origin of the recently introduced Florida population.

In contrast to *E. viridissima*, *E. dilemma* populations were genetically structured. When analyzed without *E. viridissima*, *E. dilemma* individuals clustered into three groups roughly corresponding to geography over the first two axes of a PCA (Fig. S6). These include 1) individuals from the Yucatan peninsula and northern Guatemala, 2) individuals from the allopatric southern distribution range from Tapachula in southern Mexico to northern Costa Rica and the recently established population in Florida, and 3) individuals from Veracruz (Cordoba and Los Tuxtlas). These three groups correspond to $Ed_{north}$, $Ed_{south}$, and individuals from Veracruz, which draw ancestry from multiple genetic lineages (see below), and resemble the population structure observed when analyzed together with *E. viridissima* (Fig. 2, main text). The third PC axis separated three of the four Florida individuals from all other samples (Fig. S6). One of the Florida samples clustered with the $Ed_s$ group. This suggests population structure matching both geography and allopatry – sympatry relationships with *E. viridissima*. It is unclear to what extent geography, co-occurrence with *E. viridissima*, or both might explain the observed population structure. In addition, our results indicate that the southern *E. dilemma* population ($Ed_{south}$) is the source for the recently introduced Florida population. It is thus highly likely that the introduction of *E. dilemma* to Florida was enabled by direct trade between Central American countries south of the Yucatan peninsula and Florida.

Admixture analyses support recent gene flow

The ADMIXTURE analysis revealed that several individuals drew ancestry from more than one of the three genetic lineages identified ($k$=2 and 3 most favorable cross-validation (CV) errors, mean CV error differential = 0.005; Fig. S7), therefore indicating genetic admixture or incomplete lineage sorting. For example all *E. dilemma* individuals from Veracruz (Cordoba and Los Tuxtlas, Fig. 1, main text) drew approximately 30% ancestry from $Ed_{north}$ and varying degrees from $Ed_{south}$ and $Ev$ (Fig. 2c, main text). An $f_4$-test between allopatric and sympatric groups of individuals demonstrated that the relationships between *E. dilemma* and *E. viridissima* cannot be explained by a simple bifurcating phylogeny ($f_4(Ed_{allopatric},Ed_{sympatric};Ev_{allopatric},Ev_{sympatric})$: 0.001, z : 2.7, p-value: 0.007; Table S6), supporting admixture between lineages. These observations highlight the possibility of gene flow between *E. dilemma* and *E. viridissima*.

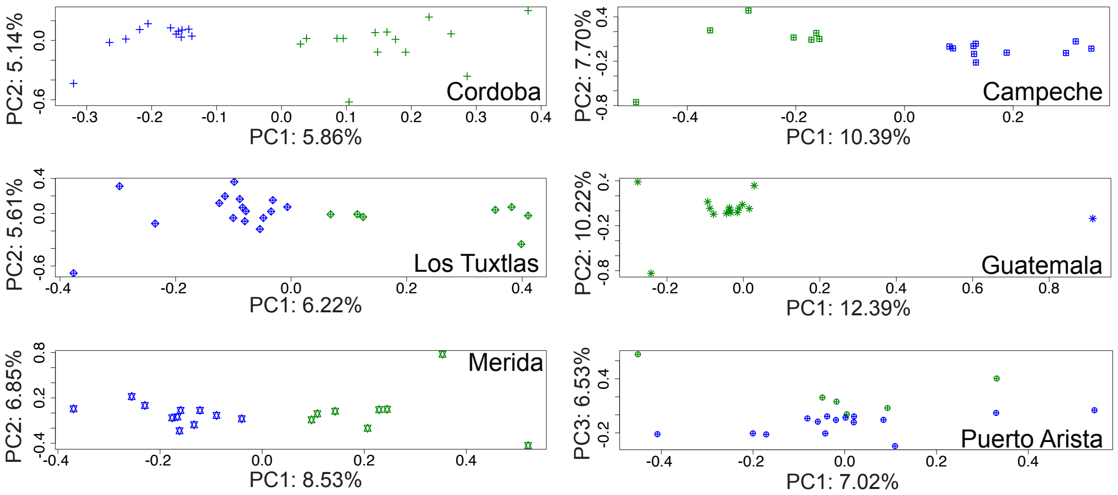

**Figure S2 PCAs of sampling sites where *E. dilemma* (green) and *E. viridissima* (blue) co-occur show that individuals cluster by species, suggesting genetic differentiation.**

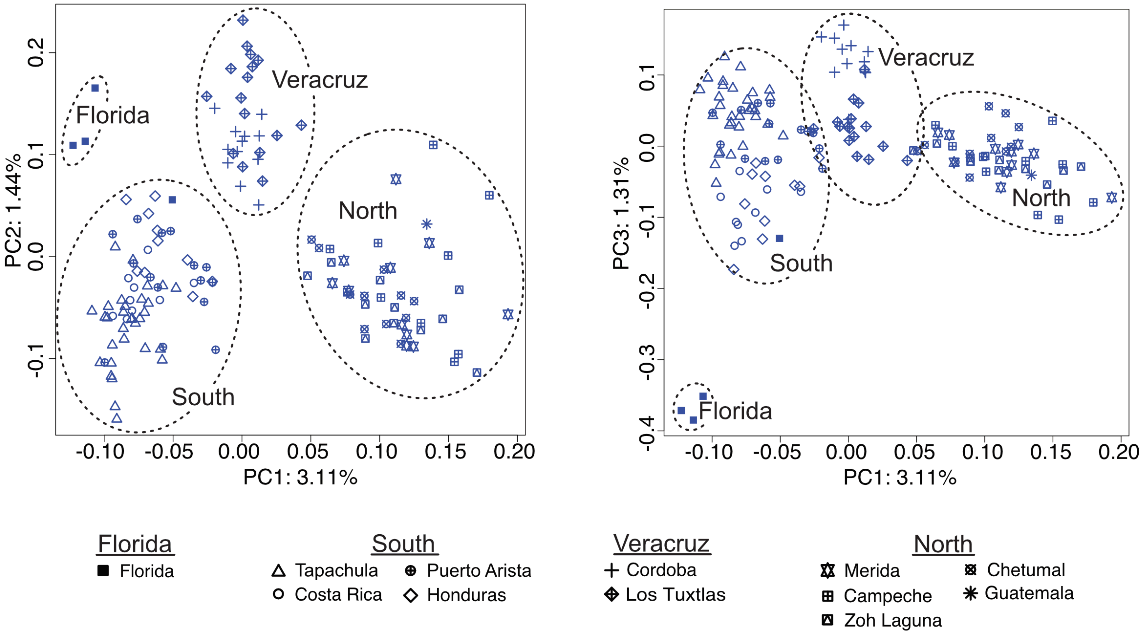

**Figure S3 Population structure within *E. dilemma*.** The first three PC axes of a PCA based on GBS data of all *E. dilemma* individuals analyzed reveal population structure roughly corresponding to geography and allopatry – sympatry with *E. dilemma*. The first PC explains about 3.11% of genetic variation, about twice as much as either PC2 or PC3. Geographic regions are indicated.

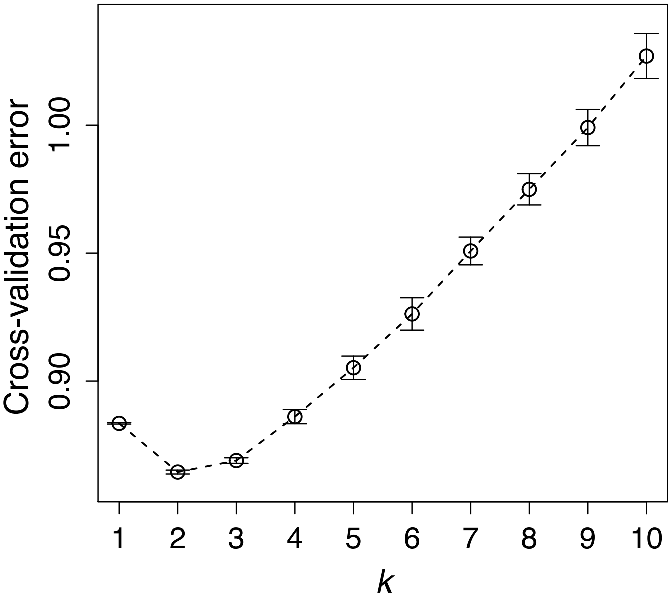

| *k* | CV error |
| --- | --- |
| 1 | 0.883 |
| 2 | 0.864 |
| 3 | 0.869 |
| 4 | 0.886 |
| 5 | 0.905 |
| 6 | 0.926 |
| 7 | 0.951 |
| 8 | 0.975 |
| 9 | 0.999 |
| 10 | 1.027 |

**Figure S4 ADMIXTURE mean cross-validation (CV) errors for *k* of 1 through 10.** Lower cross validation errors indicate higher support for a given *k* for the underlying data. *k*=2 and 3 have the most favorable cross-validation errors with a differential of less than 0.005. Error bars in the plot represent the standard deviation of the mean.

### Population genomics

Genome-wide differentiation is influenced by a common genomic landscape

Our whole genome re-sequencing approach identified 2.8 million SNPs (MAF ≥ 10%) segregating among and within species. All individuals of each genetic lineage clustered together in a PCA, supporting our GBS results (Fig. S8). Genome-wide scans revealed heterogeneous differentiation between the three genetic lineages with peaks of elevated $F_{ST}$ distributed throughout the genome (Fig. S9). Most high-differentiation regions were detected in both intra- and inter-specific pairwise comparisons (Pearson's r: 0.54 to 0.95, p=0; Fig. 3a) and therefore spanned species boundaries. Absolute sequence divergence ($D_{xy}$) and nucleotide diversity ($\pi$) were also highly correlated across genetic lineages ($D_{xy}$ r: 0.88 to 0.98, p=0; $\pi$ r: 0.95 to 0.96, p=0; $D_{xy} \sim \pi$ r: 0.82 to 0.98, p=0), suggesting that the majority of high-differentiation regions is influenced by a common genomic landscape (Poelstra *et al.* 2014; Burri *et al.* 2015; Burri 2017; Vijay *et al.* 2017) that precedes the split between *E. dilemma* and *E. viridissima*, and is likely not involved in perfume differentiation and reproductive isolation.

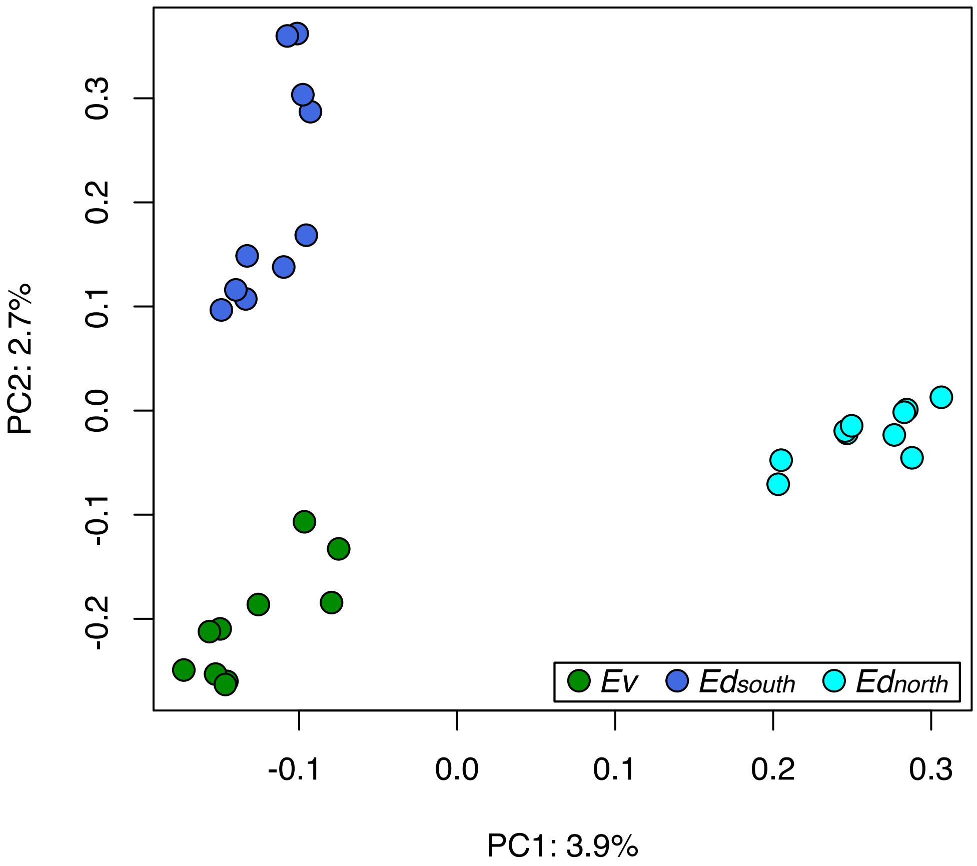

**Figure S8 PCA of whole-genome re-sequenced individuals and population relationships.** *Ev*: *E. viridissima*; $Ed_{south}$: southern *E. dilemma* population; $Ed_{north}$: northern *E. dilemma* population.

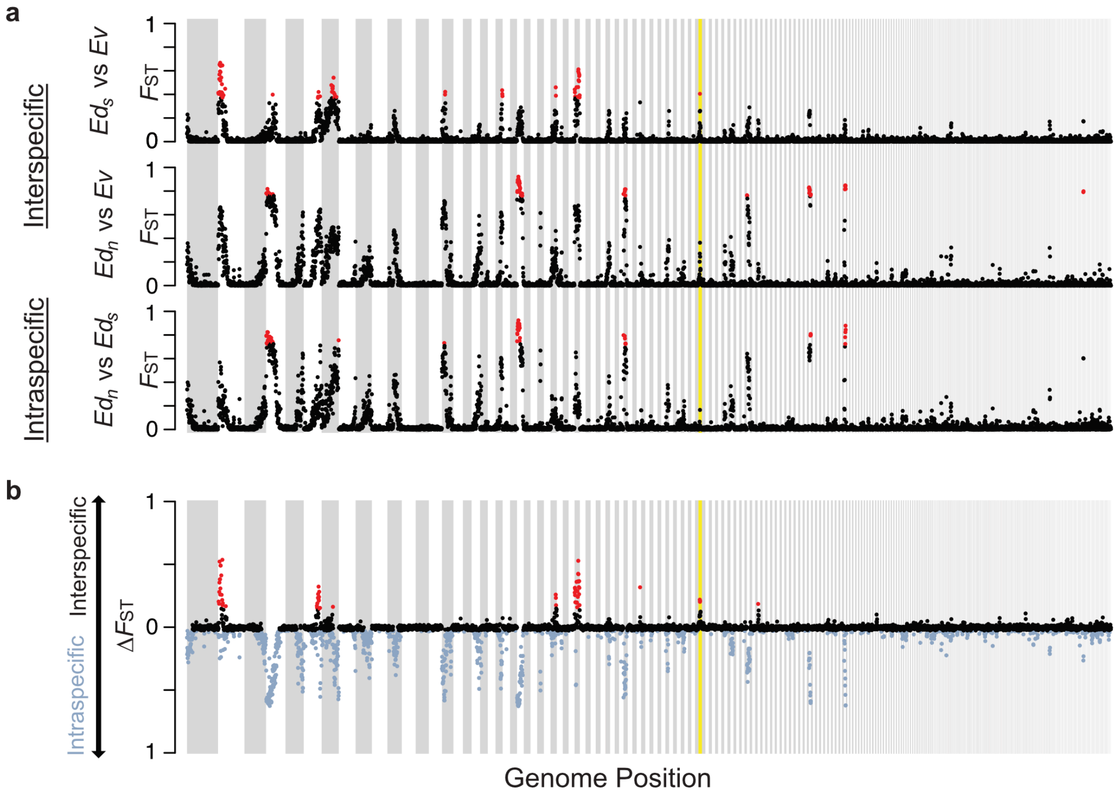

**Figure S5 Whole-genome differentiation.** (**a**) Pairwise comparisons between $Ed_{north}$, $Ed_{south}$, and $Ev$ show high heterogeneity of genetic differentiation ($F_{ST}$) throughout the genome. High-$F_{ST}$ windows (red; >99$^{th}$ percentile) clustered into peaks that were mostly shared between lineages, indicating that the genomic landscape is highly similar. (**b**) Eight regions of the genome revealed higher interspecific (black) than intraspecific (blue) differentiation ($\Delta F_{ST}$ >99$^{th}$ percentile red). The only $\Delta F_{ST}$ outlier region identified under strong divergent selection is highlighted in yellow (see main text). $Ed_n$: $Ed_{north}$, $Ed_s$: $Ed_{south}$.

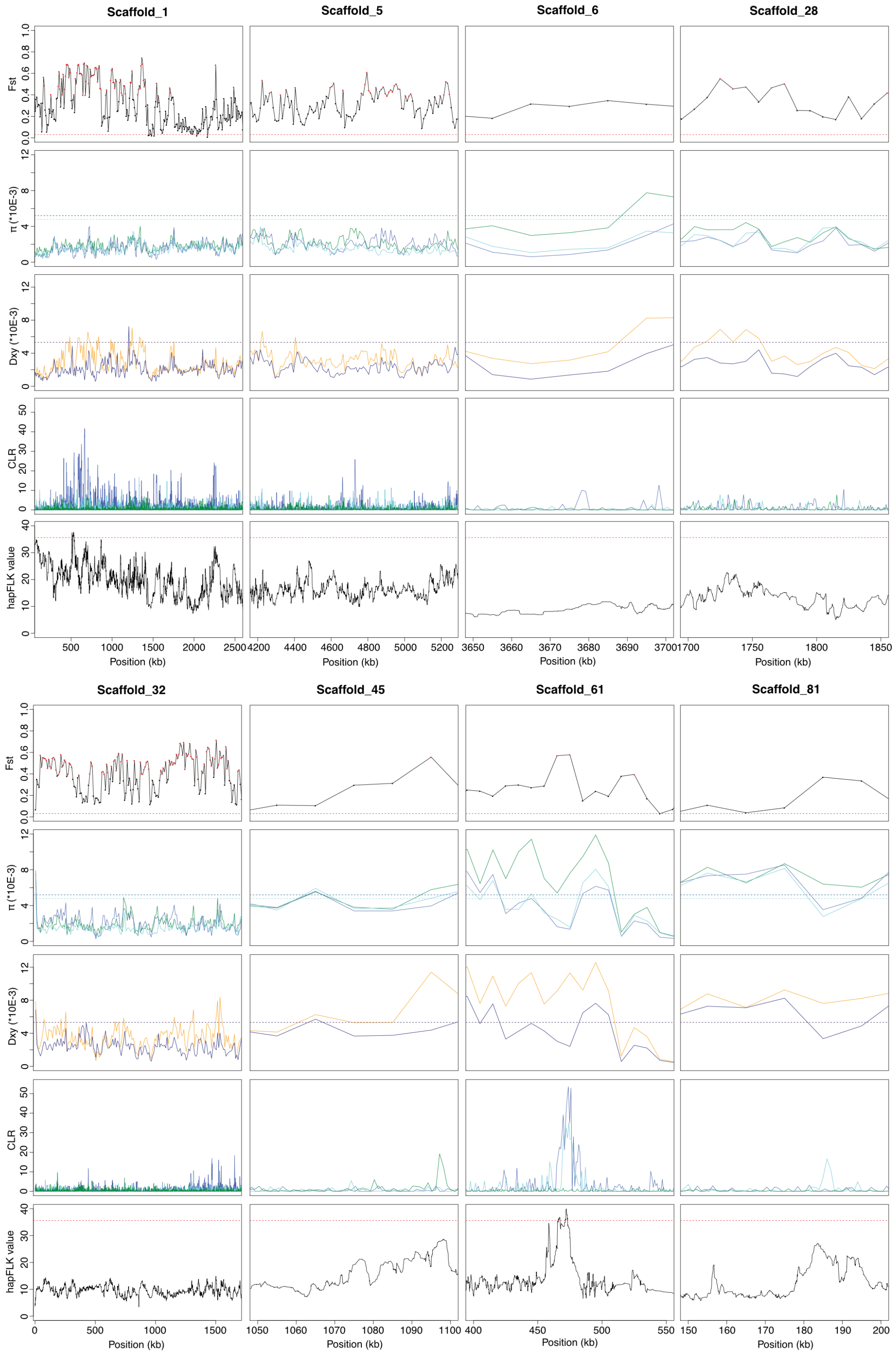

**Figure S6 Selection analysis in eight outlier peaks with elevated divergence ($\Delta F_{ST}$) between *E. dilemma* and *E. viridissima*.** Interspecific $F_{ST}$ (first row), nucleotide diversity (π, second row), absolute divergence ($D_{xy}$, third row), SweeD CLR score (fourth row), and hapFLK value (fifth row) are shown. $F_{ST}$ (first row): Red points indicate outlier 99 percentile $\Delta F_{ST}$ outlier windows (50 kb), red dotted line highlights genome-wide mean interspecific $F_{ST}$. π (second row): dark blue: $Ed_{south}$, cyan: $Ed_{north}$, green: *Ev*, dotted lines: genome-wide mean π for each lineage. $D_{xy}$ (third row): yellow (*E. dilemma* vs. *E. viridissima*), blue ($Ed_{north}$ vs. $Ed_{south}$), dotted lines: genome wide mean $D_{xy}$. Sweed CLR (fourth row): dark blue: $Ed_{south}$, cyan: $Ed_{north}$, green: *Ev*. hapFLK value (fifth row): red dotted line marks significance threshold, indicating a selective sweep. Elevated $F_{ST}$ in combination with high interspecific differential in π, elevated $D_{xy}$ between *E. dilemma* and *E. viridissima*, and lineage specific CLR outliers as well as significant hapFLK scores were observed only for the outlier region on scaffold_61 harboring OR41.

### OR41 evolutionary history

*$d_N/d_S$ test*

The likelihood of the model allowing for variation of $d_N/d_S$ among branches was significantly lower than the null model with no variation among branches (Table S11). The likelihood ratio test was 16.1 (Δ=2*((-2474.225479)-(-2482.280816))=16.1), which corresponds to a p-value <0.0001 in a $\chi^2$ distribution.

| | 40 | 43 | 44 | 56 | 70 | 80 | 86 | 99 | 118 | 129 | 131 | 162 | 167 | 217 | 275 | 305 | 317 | 403 | 407 |
|---|---|---|---|---|---|---|---|---|---|---|---|---|---|---|---|---|---|---|---|
| ***E. flammea*** | GTT | GCC | GGT | GAC | ATC | GTC | GTT | ACA | CTG | ATT | TTT | CCT | ATA | CAA | ATC | ATC | CAA | TCA | ATG |
| ***E. imperialis*** | GTT | GCC | GGT | GAC | ATC | GTC | GTT | ACA | CTG | ATT | TTT | CCT | ATA | CAA | ATC | ATC | CAA | TCG | ATG |
| ***E. viridissima*** | GTT | GCC | GGT | GAC | ATC | GTC | GTT | ACA | CTG | ATT | TTT | CCT | ATA | CAT | ATC | ATC | CAA | TCG | ATG |
| ***E. dilemma*** | CTT | GTC | GGC | GGC | CTC | ATC | ATT | ATA | GTG | GTT | GTT | CTT | GTA | GAT | GTC | GTC | CAG | ACG | ACG |

**Figure S7 Fixed substitutions in OR41 between *E. dilemma* and *E. viridissima* with respect to two outgroup species *E. flammea* and *E. imperialis*.** Substitution sites are highlighted in bold. Derived substitutions are indicated in red if non-synonymous and blue if synonymous. *E. imperialis* and *E. flammea* OR41 sequences were taken from Brand & Ramirez (2017). Numbers indicate the position of the amino acid corresponding to the nucleotide base triplet carrying a substitution.

### Supplementary Tables

**Table S1 Sampling sites**

| Bait | Latitude | Longitude | IDs | Near | Country | Sampled |
|---|---|---|---|---|---|---|
| crB04 | 9.6547333 | -85.0739333 | PB0195-PB0219 | Montezuma | Costa Rica | 4/20/15 |
| mxB01 | 18.5844833 | -95.0733833 | PB0413-PB0435 | Los Tuxtlas | Mexico | 10/3/15 |
| mxB02 | 18.5873333 | -95.07695 | PB0436-PB0457 | Los Tuxtlas | Mexico | 10/4/15 |
| mxB05 | 14.8885667 | -92.2174 | PB0471-PB0517 | Tapachula | Mexico | 10/8/15 |
| mxB06 | 15.9339 | -93.8113167 | PB0518-PB0558 | Puerto Arista | Mexico | 10/9/15 |
| mxB07 | 16.8752 | -93.4150333 | PB0559 | On the road | Mexico | 10/9/15 |
| mxB08 | 18.9155333 | -96.9823667 | PB0560-PB0588 | Córdoba | Mexico | 10/10/15 |
| mxB09 | 19.5161 | -96.9404333 | PB0589-PB0623 | Xalapa | Mexico | 10/11/15 |
| mxB12 | 20.89065 | -105.4127 | PB0675-PB0692 | Sayulita | Mexico | 10/22/15 |
| ho01 | 15.5357 | -88.3235 | PB0693-PB0708 | Cusuco | Honduras | Jun/Jul-12 |
| mxB13 | 15.6711111 | -96.5375 | SR3369-SR3393 | San Augustinillo | Mexico | 2/7/16 |
| mxB15 | 20.7889167 | -89.5905333 | PB0709-PB0769 | Merida | Mexico | 5/25/16 |
| mxB16 | 18.8662667 | -88.24675 | PB0770-PB0800 | Chetumal | Mexico | 5/26/16 |
| mxB17 | 18.6225167 | -89.3808167 | PB0801-PB0850 | Zoh-Laguna | Mexico | 5/27/16 |
| mxB18 | 19.9457667 | -90.3739833 | PB0851-PB0922 | Campeche | Mexico | 5/28/16 |
| mxB20 | 18.9611167 | -99.1091167 | PB0949-PB0980 | Tepoztlan | Mexico | 6/4/16 |
| gtB01 | 16.30504 | -89.409445 | CLY002-CLY025 | Poptún | Guatemala | Apr-16 |
| gtB02 | 14.61739 | -91.521749 | CLY026-CLY059 | Zapotitlán | Guatemala | 9/4/16 |
| usB01 | 26.228152 | -80.186781 | cd137-cd140, SR2202-SR2292 | Fern Forest | USA | 2012-2014 |

**Table S2 Individual samples.** Use of each individual in genotyping-by-sequencing (GBS), perfume phenotyping, whole-genome sequencing (WGS), and mandible morphometry is indicated. * Sampling sites correspond to Fig. 1 in the main text. ** Only males included in the morphometric analysis are indicated. *** All males with two mandibular teeth in the dataset are indicated. All others had three teeth. 1R: indicates individuals for which the right mandible was used and mirrored for morphometry. Species status is indicated (Edil: *E. dilemma*, Evir: *E. viridissima*).

| Bee ID | Bait | Sampling site* | GBS | Perfume | WGS | tridentate Mandibles** | bidentate Mandibles*** | Species |
|---|---|---|---|---|---|---|---|---|
| cd137 | usB01 | Florida | 1 | 0 | 0 | 0 | 0 | Edil |
| cd138 | usB01 | Florida | 1 | 0 | 0 | 0 | 0 | Edil |
| cd139 | usB01 | Florida | 1 | 0 | 0 | 0 | 0 | Edil |
| cd140 | usB01 | Florida | 1 | 0 | 0 | 0 | 0 | Edil |
| SR2202 | usB01 | Florida | 0 | 1 | 0 | 0 | 0 | Edil |
| SR2203 | usB01 | Florida | 0 | 1 | 0 | 0 | 0 | Edil |
| SR2204 | usB01 | Florida | 0 | 1 | 0 | 0 | 0 | Edil |
| SR2205 | usB01 | Florida | 0 | 1 | 0 | 0 | 0 | Edil |
| SR2206 | usB01 | Florida | 0 | 1 | 0 | 0 | 0 | Edil |
| SR2214 | usB01 | Florida | 0 | 1 | 0 | 0 | 0 | Edil |
| SR2215 | usB01 | Florida | 0 | 1 | 0 | 0 | 0 | Edil |
| SR2220 | usB01 | Florida | 0 | 1 | 0 | 0 | 0 | Edil |
| SR2221 | usB01 | Florida | 0 | 1 | 0 | 0 | 0 | Edil |
| SR2222 | usB01 | Florida | 0 | 1 | 0 | 0 | 0 | Edil |
| SR2225 | usB01 | Florida | 0 | 1 | 0 | 0 | 0 | Edil |
| SR2226 | usB01 | Florida | 0 | 1 | 0 | 0 | 0 | Edil |
| SR2244 | usB01 | Florida | 0 | 1 | 0 | 0 | 0 | Edil |
| SR2246 | usB01 | Florida | 0 | 1 | 0 | 0 | 0 | Edil |
| SR2247 | usB01 | Florida | 0 | 1 | 0 | 0 | 0 | Edil |
| SR2288 | usB01 | Florida | 0 | 1 | 0 | 0 | 0 | Edil |
| SR2289 | usB01 | Florida | 0 | 1 | 0 | 0 | 0 | Edil |
| SR2290 | usB01 | Florida | 0 | 1 | 0 | 0 | 0 | Edil |
| SR2291 | usB01 | Florida | 0 | 1 | 0 | 0 | 0 | Edil |
| SR2292 | usB01 | Florida | 0 | 1 | 0 | 0 | 0 | Edil |
| CLY02 | gtB01 | Guatemala | 1 | 1 | 0 | 0 | 1 | Evir |
| CLY04 | gtB01 | Guatemala | 1 | 1 | 0 | 0 | 1 | Evir |
| CLY06 | gtB01 | Guatemala | 1 | 1 | 0 | 0 | 1 | Evir |
| CLY07 | gtB01 | Guatemala | 1 | 1 | 0 | 0 | 1 | Evir |
| CLY09 | gtB01 | Guatemala | 1 | 1 | 0 | 0 | 1 | Evir |
| CLY12 | gtB01 | Guatemala | 1 | 1 | 0 | 0 | 1 | Evir |
| CLY13 | gtB01 | Guatemala | 1 | 1 | 0 | 0 | 1 | Evir |
| CLY14 | gtB01 | Guatemala | 1 | 1 | 0 | 0 | 1 | Evir |
| CLY15 | gtB01 | Guatemala | 0 | 1 | 0 | 0 | 1 | Evir |
| CLY16 | gtB01 | Guatemala | 1 | 1 | 0 | 0 | 1 | Evir |
| CLY17 | gtB01 | Guatemala | 1 | 1 | 0 | 0 | 1 | Evir |
| CLY18 | gtB01 | Guatemala | 1 | 1 | 0 | 0 | 1 | Evir |
| CLY20 | gtB01 | Guatemala | 1 | 1 | 0 | 0 | 1 | Evir |
| CLY21 | gtB01 | Guatemala | 1 | 1 | 0 | 0 | 1 | Evir |
| CLY23 | gtB01 | Guatemala | 1 | 1 | 0 | 0 | 1 | Evir |
| CLY25 | gtB01 | Guatemala | 1 | 1 | 0 | 1 | 0 | Edil |
| CLY026 | gtB02 | Tapachula | 0 | 1 | 0 | 1 | 0 | Edil |
| CLY027 | gtB02 | Tapachula | 0 | 1 | 0 | 1 | 0 | Edil |
| CLY028 | gtB02 | Tapachula | 0 | 1 | 0 | 1R | 0 | Edil |
| CLY030 | gtB02 | Tapachula | 0 | 1 | 0 | 1 | 0 | Edil |
| CLY031 | gtB02 | Tapachula | 0 | 1 | 0 | 1 | 0 | Edil |
| CLY032 | gtB02 | Tapachula | 0 | 1 | 0 | 1 | 0 | Edil |
| CLY033 | gtB02 | Tapachula | 0 | 1 | 0 | 1 | 0 | Edil |
| CLY034 | gtB02 | Tapachula | 0 | 1 | 0 | 1 | 0 | Edil |
| CLY038 | gtB02 | Tapachula | 0 | 1 | 0 | 1 | 0 | Edil |
| CLY039 | gtB02 | Tapachula | 0 | 1 | 0 | 1 | 0 | Edil |

| | | | | | | | | |
|---|---|---|---|---|---|---|---|---|
| CLY040 | gtB02 | Tapachula | 0 | 1 | 0 | 0 | 0 | Edil |
| CLY041 | gtB02 | Tapachula | 0 | 1 | 0 | 0 | 0 | Edil |
| CLY043 | gtB02 | Tapachula | 0 | 1 | 0 | 0 | 0 | Edil |
| CLY046 | gtB02 | Tapachula | 0 | 1 | 0 | 0 | 0 | Edil |
| CLY047 | gtB02 | Tapachula | 0 | 1 | 0 | 1 | 0 | Edil |
| CLY048 | gtB02 | Tapachula | 0 | 1 | 0 | 1 | 0 | Edil |
| CLY050 | gtB02 | Tapachula | 0 | 1 | 0 | 1 | 0 | Edil |
| CLY052 | gtB02 | Tapachula | 0 | 1 | 0 | 0 | 0 | Edil |
| CLY053 | gtB02 | Tapachula | 0 | 1 | 0 | 1 | 0 | Edil |
| CLY054 | gtB02 | Tapachula | 0 | 1 | 0 | 1 | 0 | Edil |
| CLY055 | gtB02 | Tapachula | 0 | 1 | 0 | 1 | 0 | Edil |
| CLY056 | gtB02 | Tapachula | 0 | 1 | 0 | 1R | 0 | Edil |
| CLY058 | gtB02 | Tapachula | 0 | 1 | 0 | 1 | 0 | Edil |
| CLY059 | gtB02 | Tapachula | 0 | 1 | 0 | 1R | 0 | Edil |
| PB0195 | crB04 | Costa Rica | 0 | 1 | 0 | 0 | 0 | Edil |
| PB0197 | crB04 | Costa Rica | 0 | 1 | 0 | 0 | 0 | Edil |
| PB0198 | crB04 | Costa Rica | 0 | 1 | 0 | 0 | 0 | Edil |
| PB0199 | crB04 | Costa Rica | 0 | 1 | 0 | 1 | 0 | Edil |
| PB0202 | crB04 | Costa Rica | 0 | 1 | 0 | 0 | 0 | Edil |
| PB0203 | crB04 | Costa Rica | 0 | 1 | 0 | 0 | 0 | Edil |
| PB0204 | crB04 | Costa Rica | 1 | 1 | 1 | 1 | 0 | Edil |
| PB0205 | crB04 | Costa Rica | 0 | 1 | 0 | 1 | 0 | Edil |
| PB0206 | crB04 | Costa Rica | 1 | 1 | 1 | 1 | 0 | Edil |
| PB0207 | crB04 | Costa Rica | 0 | 1 | 0 | 0 | 0 | Edil |
| PB0208 | crB04 | Costa Rica | 0 | 1 | 0 | 0 | 0 | Edil |
| PB0209 | crB04 | Costa Rica | 1 | 1 | 1 | 1 | 0 | Edil |
| PB0210 | crB04 | Costa Rica | 1 | 1 | 1 | 1 | 0 | Edil |
| PB0211 | crB04 | Costa Rica | 1 | 1 | 0 | 1 | 0 | Edil |
| PB0212 | crB04 | Costa Rica | 0 | 1 | 0 | 1 | 0 | Edil |
| PB0214 | crB04 | Costa Rica | 0 | 1 | 0 | 0 | 0 | Edil |
| PB0215 | crB04 | Costa Rica | 1 | 1 | 0 | 1 | 0 | Edil |
| PB0216 | crB04 | Costa Rica | 1 | 1 | 1 | 1 | 0 | Edil |
| PB0217 | crB04 | Costa Rica | 1 | 1 | 0 | 1 | 0 | Edil |
| PB0219 | crB04 | Costa Rica | 1 | 1 | 0 | 1 | 0 | Edil |
| PB0413 | mxB01 | Los Tuxtlas | 1 | 1 | 0 | 0 | 1 | Evir |
| PB0415 | mxB01 | Los Tuxtlas | 1 | 1 | 0 | 1 | 0 | Edil |
| PB0417 | mxB01 | Los Tuxtlas | 1 | 1 | 0 | 1 | 0 | Evir |
| PB0419 | mxB01 | Los Tuxtlas | 1 | 1 | 0 | 0 | 1 | Evir |
| PB0420 | mxB01 | Los Tuxtlas | 1 | 1 | 0 | 0 | 1 | Evir |
| PB0421 | mxB01 | Los Tuxtlas | 1 | 1 | 1 | 0 | 1 | Evir |
| PB0424 | mxB01 | Los Tuxtlas | 1 | 1 | 0 | 1 | 0 | Edil |
| PB0427 | mxB01 | Los Tuxtlas | 1 | 1 | 0 | 1 | 0 | Edil |
| PB0429 | mxB01 | Los Tuxtlas | 1 | 1 | 0 | 1 | 0 | Edil |
| PB0434 | mxB01 | Los Tuxtlas | 1 | 1 | 0 | 1 | 0 | Edil |
| PB0435 | mxB01 | Los Tuxtlas | 1 | 1 | 0 | 1 | 0 | Edil |
| PB0437 | mxB02 | Los Tuxtlas | 1 | 1 | 0 | 1 | 0 | Edil |
| PB0441 | mxB02 | Los Tuxtlas | 1 | 1 | 0 | 1 | 0 | Edil |
| PB0442 | mxB02 | Los Tuxtlas | 1 | 1 | 0 | 1 | 0 | Edil |
| PB0443 | mxB02 | Los Tuxtlas | 1 | 1 | 0 | 1 | 0 | Edil |
| PB0444 | mxB02 | Los Tuxtlas | 1 | 1 | 0 | 1 | 0 | Edil |
| PB0445 | mxB02 | Los Tuxtlas | 1 | 1 | 0 | 0 | 1 | Evir |
| PB0446 | mxB02 | Los Tuxtlas | 1 | 1 | 0 | 0 | 1 | Evir |
| PB0449 | mxB02 | Los Tuxtlas | 1 | 1 | 0 | 0 | 1 | Evir |
| PB0450 | mxB02 | Los Tuxtlas | 1 | 1 | 0 | 1 | 0 | Edil |
| PB0451 | mxB02 | Los Tuxtlas | 1 | 1 | 0 | 1 | 0 | Edil |
| PB0455 | mxB02 | Los Tuxtlas | 1 | 1 | 0 | 1 | 0 | Edil |
| PB0457 | mxB02 | Los Tuxtlas | 1 | 1 | 0 | 1 | 0 | Edil |
| PB0471 | mxB05 | Tapachula | 1 | 1 | 1 | 1 | 0 | Edil |
| PB0472 | mxB05 | Tapachula | 1 | 1 | 0 | 1 | 0 | Edil |

| | | | | | | | | |
|---|---|---|---|---|---|---|---|---|
| PB0473 | mxB05 | Tapachula | 1 | 1 | 0 | 1 | 0 | Edil |
| PB0474 | mxB05 | Tapachula | 1 | 1 | 0 | 1 | 0 | Edil |
| PB0476 | mxB05 | Tapachula | 1 | 1 | 0 | 1 | 0 | Edil |
| PB0477 | mxB05 | Tapachula | 1 | 1 | 0 | 1R | 0 | Edil |
| PB0478 | mxB05 | Tapachula | 0 | 0 | 0 | 1 | 0 | Edil |
| PB0479 | mxB05 | Tapachula | 0 | 0 | 0 | 1 | 0 | Edil |
| PB0480 | mxB05 | Tapachula | 1 | 1 | 0 | 1 | 0 | Edil |
| PB0481 | mxB05 | Tapachula | 1 | 1 | 0 | 1 | 0 | Edil |
| PB0482 | mxB05 | Tapachula | 1 | 1 | 1 | 1R | 0 | Edil |
| PB0483 | mxB05 | Tapachula | 1 | 1 | 0 | 1 | 0 | Edil |
| PB0484 | mxB05 | Tapachula | 1 | 1 | 0 | 1 | 0 | Edil |
| PB0485 | mxB05 | Tapachula | 1 | 1 | 1 | 1 | 0 | Edil |
| PB0486 | mxB05 | Tapachula | 0 | 0 | 0 | 1 | 0 | Edil |
| PB0487 | mxB05 | Tapachula | 1 | 1 | 0 | 1 | 0 | Edil |
| PB0488 | mxB05 | Tapachula | 0 | 0 | 0 | 1 | 0 | Edil |
| PB0489 | mxB05 | Tapachula | 1 | 1 | 0 | 1 | 0 | Edil |
| PB0490 | mxB05 | Tapachula | 1 | 1 | 1 | 1 | 0 | Edil |
| PB0491 | mxB05 | Tapachula | 1 | 1 | 0 | 1R | 0 | Edil |
| PB0492 | mxB05 | Tapachula | 1 | 1 | 0 | 1 | 0 | Edil |
| PB0494 | mxB05 | Tapachula | 1 | 1 | 0 | 0 | 0 | Edil |
| PB0496 | mxB05 | Tapachula | 1 | 1 | 0 | 1 | 0 | Edil |
| PB0497 | mxB05 | Tapachula | 1 | 1 | 1 | 1 | 0 | Edil |
| PB0498 | mxB05 | Tapachula | 0 | 0 | 0 | 1 | 0 | Edil |
| PB0499 | mxB05 | Tapachula | 1 | 1 | 0 | 1 | 0 | Edil |
| PB0500 | mxB05 | Tapachula | 1 | 1 | 0 | 1 | 0 | Edil |
| PB0501 | mxB05 | Tapachula | 1 | 1 | 0 | 1R | 0 | Edil |
| PB0502 | mxB05 | Tapachula | 0 | 1 | 0 | 1R | 0 | Edil |
| PB0503 | mxB05 | Tapachula | 1 | 1 | 0 | 1 | 0 | Edil |
| PB0505 | mxB05 | Tapachula | 1 | 1 | 0 | 1 | 0 | Edil |
| PB0506 | mxB05 | Tapachula | 1 | 1 | 0 | 1 | 0 | Edil |
| PB0507 | mxB05 | Tapachula | 1 | 1 | 0 | 1 | 0 | Edil |
| PB0508 | mxB05 | Tapachula | 0 | 0 | 0 | 1 | 0 | Edil |
| PB0509 | mxB05 | Tapachula | 1 | 1 | 0 | 1R | 0 | Edil |
| PB0510 | mxB05 | Tapachula | 0 | 0 | 0 | 1 | 0 | Edil |
| PB0511 | mxB05 | Tapachula | 0 | 0 | 0 | 1 | 0 | Edil |
| PB0512 | mxB05 | Tapachula | 1 | 1 | 0 | 1 | 0 | Edil |
| PB0513 | mxB05 | Tapachula | 1 | 1 | 0 | 1R | 0 | Edil |
| PB0514 | mxB05 | Tapachula | 0 | 0 | 0 | 1 | 0 | Edil |
| PB0515 | mxB05 | Tapachula | 0 | 0 | 0 | 1 | 0 | Edil |
| PB0516 | mxB05 | Tapachula | 1 | 1 | 0 | 1 | 0 | Edil |
| PB0517 | mxB05 | Tapachula | 0 | 0 | 0 | 1 | 0 | Edil |
| PB0518 | mxB06 | Puerto Arista | 0 | 1 | 0 | 0 | 0 | Edil |
| PB0519 | mxB06 | Puerto Arista | 1 | 1 | 0 | 0 | 0 | Edil |
| PB0520 | mxB06 | Puerto Arista | 1 | 1 | 0 | 0 | 0 | Edil |
| PB0521 | mxB06 | Puerto Arista | 1 | 1 | 0 | 0 | 0 | Edil |
| PB0523 | mxB06 | Puerto Arista | 0 | 1 | 0 | 0 | 0 | Edil |
| PB0525 | mxB06 | Puerto Arista | 0 | 1 | 0 | 0 | 0 | Edil |
| PB0526 | mxB06 | Puerto Arista | 1 | 1 | 0 | 1 | 0 | Edil |
| PB0527 | mxB06 | Puerto Arista | 1 | 1 | 0 | 1 | 0 | Edil |
| PB0528 | mxB06 | Puerto Arista | 1 | 1 | 0 | 0 | 1 | Evir |
| PB0529 | mxB06 | Puerto Arista | 0 | 1 | 0 | 0 | 0 | Edil |
| PB0530 | mxB06 | Puerto Arista | 1 | 1 | 0 | 0 | 0 | Edil |
| PB0533 | mxB06 | Puerto Arista | 1 | 1 | 0 | 0 | 1 | Evir |
| PB0534 | mxB06 | Puerto Arista | 1 | 1 | 0 | 0 | 0 | Evir |
| PB0535 | mxB06 | Puerto Arista | 0 | 1 | 0 | 0 | 0 | Edil |
| PB0536 | mxB06 | Puerto Arista | 0 | 1 | 0 | 1 | 0 | Edil |
| PB0537 | mxB06 | Puerto Arista | 1 | 1 | 0 | 0 | 1 | Evir |
| PB0538 | mxB06 | Puerto Arista | 0 | 1 | 0 | 0 | 0 | Edil |
| PB0539 | mxB06 | Puerto Arista | 0 | 1 | 0 | 0 | 0 | Edil |

| | | | | | | | | |
|---|---|---|---|---|---|---|---|---|
| PB0540 | mxB06 | Puerto Arista | 1 | 1 | 0 | 1 | 0 | Evir |
| PB0541 | mxB06 | Puerto Arista | 1 | 1 | 0 | 0 | 0 | Edil |
| PB0542 | mxB06 | Puerto Arista | 0 | 1 | 0 | 1 | 0 | Edil |
| PB0543 | mxB06 | Puerto Arista | 1 | 1 | 0 | 1 | 0 | Edil |
| PB0544 | mxB06 | Puerto Arista | 0 | 1 | 0 | 1 | 0 | Edil |
| PB0545 | mxB06 | Puerto Arista | 1 | 1 | 0 | 1 | 0 | Evir |
| PB0546 | mxB06 | Puerto Arista | 1 | 1 | 0 | 0 | 1 | Evir |
| PB0547 | mxB06 | Puerto Arista | 1 | 1 | 0 | 1 | 0 | Edil |
| PB0548 | mxB06 | Puerto Arista | 1 | 1 | 0 | 0 | 1 | Evir |
| PB0549 | mxB06 | Puerto Arista | 1 | 1 | 0 | 1 | 0 | Edil |
| PB0550 | mxB06 | Puerto Arista | 0 | 1 | 0 | 0 | 0 | Edil |
| PB0551 | mxB06 | Puerto Arista | 0 | 1 | 0 | 0 | 0 | Edil |
| PB0554 | mxB06 | Puerto Arista | 1 | 1 | 0 | 0 | 0 | Edil |
| PB0555 | mxB06 | Puerto Arista | 0 | 1 | 0 | 0 | 0 | Edil |
| PB0556 | mxB06 | Puerto Arista | 0 | 1 | 0 | 0 | 0 | Edil |
| PB0557 | mxB06 | Puerto Arista | 0 | 1 | 0 | 0 | 0 | Edil |
| PB0558 | mxB06 | Puerto Arista | 1 | 1 | 0 | 0 | 1 | Evir |
| PB0559 | mxB07 | Los Tuxtlas | 1 | 0 | 0 | 0 | 0 | Edil |
| PB0560 | mxB08 | Córdoba | 1 | 1 | 0 | 0 | 0 | Evir |
| PB0561 | mxB08 | Córdoba | 1 | 1 | 0 | 0 | 0 | Evir |
| PB0562 | mxB08 | Córdoba | 1 | 1 | 0 | 0 | 0 | Evir |
| PB0563 | mxB08 | Córdoba | 1 | 1 | 0 | 0 | 0 | Evir |
| PB0564 | mxB08 | Córdoba | 1 | 1 | 0 | 0 | 0 | Evir |
| PB0565 | mxB08 | Córdoba | 1 | 1 | 0 | 0 | 0 | Evir |
| PB0566 | mxB08 | Córdoba | 1 | 1 | 0 | 1 | 0 | Edil |
| PB0567 | mxB08 | Córdoba | 1 | 1 | 0 | 1 | 0 | Edil |
| PB0568 | mxB08 | Córdoba | 1 | 1 | 0 | 1 | 0 | Edil |
| PB0571 | mxB08 | Córdoba | 0 | 1 | 0 | 1R | 0 | Edil |
| PB0576 | mxB08 | Córdoba | 1 | 1 | 0 | 1 | 0 | Edil |
| PB0579 | mxB08 | Córdoba | 1 | 1 | 0 | 1 | 0 | Edil |
| PB0589 | mxB09 | Córdoba | 1 | 1 | 0 | 1R | 0 | Evir |
| PB0590 | mxB09 | Córdoba | 1 | 1 | 0 | 0 | 0 | Evir |
| PB0591 | mxB08 | Córdoba | 0 | 1 | 0 | 0 | 0 | Evir |
| PB0592 | mxB09 | Córdoba | 1 | 1 | 0 | 1 | 0 | Evir |
| PB0593 | mxB09 | Córdoba | 1 | 1 | 0 | 0 | 0 | Evir |
| PB0594 | mxB09 | Córdoba | 1 | 1 | 0 | 1 | 0 | Edil |
| PB0595 | mxB09 | Córdoba | 1 | 1 | 0 | 1 | 0 | Edil |
| PB0596 | mxB09 | Córdoba | 1 | 1 | 0 | 0 | 0 | Evir |
| PB0597 | mxB09 | Córdoba | 1 | 1 | 0 | 0 | 0 | Evir |
| PB0598 | mxB09 | Córdoba | 1 | 1 | 0 | 1R | 0 | Edil |
| PB0599 | mxB09 | Córdoba | 1 | 1 | 0 | 0 | 0 | Evir |
| PB0601 | mxB09 | Córdoba | 1 | 1 | 0 | 1 | 0 | Evir |
| PB0602 | mxB09 | Córdoba | 1 | 1 | 0 | 1R | 0 | Evir |
| PB0614 | mxB09 | Córdoba | 1 | 1 | 0 | 1 | 0 | Edil |
| PB0616 | mxB09 | Córdoba | 1 | 1 | 0 | 1 | 0 | Edil |
| PB0618 | mxB09 | Córdoba | 1 | 1 | 0 | 1R | 0 | Edil |
| PB0624 | mxB12 | Sayulita | 0 | 1 | 0 | 0 | 1 | Evir |
| PB0625 | mxB12 | Sayulita | 0 | 1 | 0 | 0 | 1 | Evir |
| PB0626 | mxB12 | Sayulita | 0 | 1 | 0 | 0 | 1 | Evir |
| PB0627 | mxB12 | Sayulita | 0 | 1 | 0 | 0 | 1 | Evir |
| PB0628 | mxB12 | Sayulita | 0 | 1 | 0 | 0 | 1 | Evir |
| PB0629 | mxB12 | Sayulita | 0 | 1 | 0 | 0 | 1 | Evir |
| PB0630 | mxB12 | Sayulita | 0 | 1 | 0 | 0 | 1 | Evir |
| PB0631 | mxB12 | Sayulita | 0 | 1 | 0 | 0 | 1 | Evir |
| PB0632 | mxB12 | Sayulita | 0 | 1 | 0 | 0 | 1 | Evir |
| PB0633 | mxB12 | Sayulita | 0 | 1 | 0 | 0 | 1 | Evir |
| PB0634 | mxB12 | Sayulita | 0 | 1 | 0 | 0 | 1 | Evir |
| PB0635 | mxB12 | Sayulita | 0 | 1 | 0 | 0 | 1 | Evir |
| PB0636 | mxB12 | Sayulita | 0 | 1 | 0 | 0 | 1 | Evir |

| | | | | | | | | |
|---|---|---|---|---|---|---|---|---|
| PB0637 | mxB12 | Sayulita | 0 | 1 | 0 | 0 | 1 | Evir |
| PB0638 | mxB12 | Sayulita | 0 | 1 | 0 | 0 | 1 | Evir |
| PB0639 | mxB12 | Sayulita | 0 | 1 | 0 | 0 | 1 | Evir |
| PB0640 | mxB12 | Sayulita | 0 | 1 | 0 | 0 | 1 | Evir |
| PB0641 | mxB12 | Sayulita | 0 | 1 | 0 | 0 | 1 | Evir |
| PB0642 | mxB12 | Sayulita | 0 | 1 | 0 | 0 | 1 | Evir |
| PB0643 | mxB12 | Sayulita | 0 | 1 | 0 | 0 | 1 | Evir |
| PB0675 | mxB12 | Sayulita | 1 | 1 | 0 | 0 | 1 | Evir |
| PB0676 | mxB12 | Sayulita | 1 | 1 | 0 | 0 | 1 | Evir |
| PB0677 | mxB12 | Sayulita | 1 | 1 | 0 | 0 | 1 | Evir |
| PB0678 | mxB12 | Sayulita | 1 | 1 | 0 | 0 | 1 | Evir |
| PB0679 | mxB12 | Sayulita | 1 | 1 | 1 | 0 | 1 | Evir |
| PB0680 | mxB12 | Sayulita | 1 | 1 | 0 | 0 | 1 | Evir |
| PB0681 | mxB12 | Sayulita | 1 | 1 | 1 | 0 | 1 | Evir |
| PB0682 | mxB12 | Sayulita | 1 | 1 | 0 | 0 | 1 | Evir |
| PB0683 | mxB12 | Sayulita | 1 | 1 | 0 | 0 | 1 | Evir |
| PB0684 | mxB12 | Sayulita | 1 | 1 | 0 | 0 | 1 | Evir |
| PB0685 | mxB12 | Sayulita | 1 | 1 | 0 | 0 | 1 | Evir |
| PB0686 | mxB12 | Sayulita | 1 | 1 | 0 | 0 | 1 | Evir |
| PB0687 | mxB12 | Sayulita | 0 | 1 | 0 | 0 | 1 | Evir |
| PB0688 | mxB12 | Sayulita | 0 | 1 | 0 | 0 | 1 | Evir |
| PB0689 | mxB12 | Sayulita | 0 | 1 | 0 | 0 | 1 | Evir |
| PB0690 | mxB12 | Sayulita | 0 | 1 | 0 | 0 | 1 | Evir |
| PB0691 | mxB12 | Sayulita | 0 | 1 | 0 | 0 | 1 | Evir |
| PB0692 | mxB12 | Sayulita | 0 | 1 | 0 | 0 | 1 | Evir |
| PB0697 | ho01 | Honduras | 1 | 0 | 0 | 0 | 0 | Edil |
| PB0698 | ho01 | Honduras | 1 | 0 | 0 | 0 | 0 | Edil |
| PB0699 | ho01 | Honduras | 1 | 0 | 0 | 0 | 0 | Edil |
| PB0700 | ho01 | Honduras | 1 | 0 | 0 | 0 | 0 | Edil |
| PB0701 | ho01 | Honduras | 1 | 0 | 0 | 0 | 0 | Edil |
| PB0702 | ho01 | Honduras | 1 | 0 | 0 | 0 | 0 | Edil |
| PB0705 | ho01 | Honduras | 1 | 0 | 0 | 0 | 0 | Edil |
| PB0706 | ho01 | Honduras | 1 | 0 | 0 | 0 | 0 | Edil |
| PB0707 | ho01 | Honduras | 1 | 0 | 0 | 0 | 0 | Edil |
| PB0708 | ho01 | Honduras | 1 | 0 | 0 | 0 | 0 | Edil |
| PB0709 | mxB15 | Mérida | 1 | 1 | 1 | 1 | 0 | Edil |
| PB0710 | mxB15 | Mérida | 1 | 1 | 0 | 1 | 0 | Edil |
| PB0711 | mxB15 | Mérida | 0 | 1 | 0 | 0 | 0 | Edil |
| PB0713 | mxB15 | Mérida | 1 | 1 | 0 | 0 | 0 | Edil |
| PB0716 | mxB15 | Mérida | 1 | 1 | 0 | 0 | 1 | Evir |
| PB0718 | mxB15 | Mérida | 1 | 1 | 1 | 1 | 0 | Edil |
| PB0719 | mxB15 | Mérida | 0 | 1 | 0 | 0 | 1 | Evir |
| PB0721 | mxB15 | Mérida | 1 | 1 | 0 | 0 | 0 | Edil |
| PB0722 | mxB15 | Mérida | 0 | 1 | 0 | 0 | 1 | Evir |
| PB0723 | mxB15 | Mérida | 0 | 1 | 0 | 1 | 0 | Evir |
| PB0729 | mxB15 | Mérida | 1 | 1 | 1 | 0 | 1 | Evir |
| PB0736 | mxB15 | Mérida | 1 | 1 | 0 | 1 | 0 | Edil |
| PB0738 | mxB15 | Mérida | 1 | 1 | 1 | 0 | 1 | Evir |
| PB0740 | mxB15 | Mérida | 1 | 1 | 1 | 0 | 0 | Edil |
| PB0743 | mxB15 | Mérida | 1 | 1 | 0 | 0 | 1 | Evir |
| PB0745 | mxB15 | Mérida | 0 | 1 | 0 | 0 | 1 | Evir |
| PB0749 | mxB15 | Mérida | 1 | 1 | 1 | 1 | 0 | Edil |
| PB0754 | mxB15 | Mérida | 1 | 1 | 1 | 0 | 0 | Edil |
| PB0758 | mxB15 | Mérida | 1 | 1 | 0 | 0 | 1 | Evir |
| PB0760 | mxB15 | Mérida | 1 | 1 | 0 | 0 | 1 | Evir |
| PB0761 | mxB15 | Mérida | 1 | 1 | 0 | 0 | 1 | Evir |
| PB0763 | mxB15 | Mérida | 1 | 1 | 0 | 0 | 0 | Edil |
| PB0764 | mxB15 | Mérida | 0 | 1 | 0 | 0 | 1 | Evir |
| PB0766 | mxB15 | Mérida | 1 | 1 | 0 | 0 | 1 | Evir |

| | | | | | | | | |
|---|---|---|---|---|---|---|---|---|
| PB0769 | mxB15 | Mérida | 1 | 1 | 0 | 0 | 1 | Evir |
| PB0770 | mxB16 | Chetumal | 0 | 1 | 0 | 0 | 0 | Edil |
| PB0771 | mxB16 | Chetumal | 0 | 1 | 0 | 0 | 0 | Edil |
| PB0772 | mxB16 | Chetumal | 1 | 1 | 0 | 0 | 0 | Edil |
| PB0773 | mxB16 | Chetumal | 1 | 1 | 0 | 0 | 0 | Edil |
| PB0774 | mxB16 | Chetumal | 0 | 1 | 0 | 1 | 0 | Edil |
| PB0775 | mxB16 | Chetumal | 1 | 1 | 0 | 1 | 0 | Edil |
| PB0776 | mxB16 | Chetumal | 0 | 1 | 0 | 0 | 0 | Edil |
| PB0777 | mxB16 | Chetumal | 0 | 1 | 0 | 1 | 0 | Edil |
| PB0778 | mxB16 | Chetumal | 1 | 1 | 0 | 1R | 0 | Edil |
| PB0779 | mxB16 | Chetumal | 0 | 1 | 0 | 1 | 0 | Edil |
| PB0780 | mxB16 | Chetumal | 1 | 1 | 0 | 1 | 0 | Edil |
| PB0781 | mxB16 | Chetumal | 1 | 1 | 0 | 1 | 0 | Edil |
| PB0782 | mxB16 | Chetumal | 0 | 1 | 0 | 0 | 0 | Edil |
| PB0783 | mxB16 | Chetumal | 0 | 1 | 0 | 0 | 0 | Edil |
| PB0784 | mxB16 | Chetumal | 0 | 1 | 0 | 0 | 0 | Edil |
| PB0785 | mxB16 | Chetumal | 0 | 1 | 0 | 0 | 0 | Edil |
| PB0786 | mxB16 | Chetumal | 1 | 1 | 0 | 1 | 0 | Edil |
| PB0787 | mxB16 | Chetumal | 0 | 1 | 0 | 1 | 0 | Edil |
| PB0788 | mxB16 | Chetumal | 0 | 1 | 0 | 0 | 0 | Edil |
| PB0789 | mxB16 | Chetumal | 0 | 1 | 0 | 1 | 0 | Edil |
| PB0790 | mxB16 | Chetumal | 0 | 1 | 0 | 1 | 0 | Edil |
| PB0791 | mxB16 | Chetumal | 0 | 1 | 0 | 0 | 0 | Edil |
| PB0792 | mxB16 | Chetumal | 1 | 1 | 0 | 1 | 0 | Edil |
| PB0793 | mxB16 | Chetumal | 0 | 1 | 0 | 1 | 0 | Edil |
| PB0794 | mxB16 | Chetumal | 0 | 1 | 0 | 0 | 0 | Edil |
| PB0795 | mxB16 | Chetumal | 0 | 1 | 0 | 1 | 0 | Edil |
| PB0796 | mxB16 | Chetumal | 0 | 1 | 0 | 0 | 0 | Edil |
| PB0797 | mxB16 | Chetumal | 1 | 1 | 0 | 1 | 0 | Edil |
| PB0798 | mxB16 | Chetumal | 0 | 1 | 0 | 1 | 0 | Edil |
| PB0799 | mxB16 | Chetumal | 1 | 1 | 0 | 1 | 0 | Edil |
| PB0800 | mxB16 | Chetumal | 1 | 1 | 0 | 1 | 0 | Edil |
| PB0801 | mxB17 | Zoh-laguna | 1 | 1 | 0 | 1 | 0 | Edil |
| PB0802 | mxB17 | Zoh-laguna | 1 | 1 | 0 | 1 | 0 | Edil |
| PB0803 | mxB17 | Zoh-laguna | 0 | 1 | 0 | 1 | 0 | Edil |
| PB0804 | mxB17 | Zoh-laguna | 0 | 1 | 0 | 0 | 0 | Edil |
| PB0805 | mxB17 | Zoh-laguna | 0 | 1 | 0 | 1 | 0 | Edil |
| PB0806 | mxB17 | Zoh-laguna | 0 | 1 | 0 | 0 | 0 | Edil |
| PB0807 | mxB17 | Zoh-laguna | 1 | 1 | 0 | 1 | 0 | Edil |
| PB0808 | mxB17 | Zoh-laguna | 1 | 1 | 0 | 1 | 0 | Edil |
| PB0809 | mxB17 | Zoh-laguna | 0 | 1 | 0 | 1 | 0 | Edil |
| PB0810 | mxB17 | Zoh-laguna | 1 | 1 | 0 | 1 | 0 | Evir |
| PB0811 | mxB17 | Zoh-laguna | 0 | 1 | 0 | 1 | 0 | Evir |
| PB0812 | mxB17 | Zoh-laguna | 0 | 1 | 0 | 1 | 0 | Edil |
| PB0813 | mxB17 | Zoh-laguna | 0 | 1 | 0 | 0 | 0 | Edil |
| PB0814 | mxB17 | Zoh-laguna | 0 | 1 | 0 | 1 | 0 | Edil |
| PB0815 | mxB17 | Zoh-laguna | 1 | 1 | 0 | 1 | 0 | Edil |
| PB0816 | mxB17 | Zoh-laguna | 1 | 1 | 0 | 1 | 0 | Edil |
| PB0817 | mxB17 | Zoh-laguna | 1 | 1 | 0 | 1 | 0 | Edil |
| PB0818 | mxB17 | Zoh-laguna | 1 | 1 | 0 | 1 | 0 | Edil |
| PB0819 | mxB17 | Zoh-laguna | 0 | 1 | 0 | 0 | 0 | Edil |
| PB0820 | mxB17 | Zoh-laguna | 0 | 1 | 0 | 1 | 0 | Edil |
| PB0837 | mxB17 | Zoh-laguna | 1 | 0 | 0 | 1 | 0 | Edil |
| PB0844 | mxB17 | Zoh-laguna | 1 | 0 | 0 | 1R | 0 | Edil |
| PB0856 | mxB18 | Campeche | 1 | 1 | 1 | 1 | 0 | Edil |
| PB0858 | mxB18 | Campeche | 1 | 1 | 0 | 1 | 0 | Evir |
| PB0862 | mxB18 | Campeche | 1 | 1 | 1 | 1 | 0 | Edil |
| PB0863 | mxB18 | Campeche | 1 | 1 | 0 | 0 | 1 | Evir |
| PB0864 | mxB18 | Campeche | 1 | 1 | 1 | 1 | 0 | Edil |

| | | | | | | | | |
|---|---|---|---|---|---|---|---|---|
| PB0867 | mxB18 | Campeche | 1 | 1 | 0 | 1 | 0 | Edil |
| PB0869 | mxB18 | Campeche | 0 | 1 | 0 | 0 | 0 | Edil |
| PB0877 | mxB18 | Campeche | 1 | 1 | 0 | 1 | 0 | Evir |
| PB0878 | mxB18 | Campeche | 1 | 1 | 0 | 0 | 1 | Evir |
| PB0887 | mxB18 | Campeche | 1 | 1 | 0 | 1 | 0 | Edil |
| PB0891 | mxB18 | Campeche | 1 | 1 | 1 | 0 | 1 | Evir |
| PB0900 | mxB18 | Campeche | 1 | 1 | 1 | 1 | 0 | Edil |
| PB0904 | mxB18 | Campeche | 0 | 1 | 0 | 0 | 1 | Evir |
| PB0906 | mxB18 | Campeche | 0 | 1 | 0 | 0 | 1 | Evir |
| PB0909 | mxB18 | Campeche | 1 | 1 | 1 | 0 | 1 | Evir |
| PB0911 | mxB18 | Campeche | 1 | 1 | 0 | 1 | 0 | Edil |
| PB0913 | mxB18 | Campeche | 1 | 1 | 0 | 1 | 0 | Edil |
| PB0915 | mxB18 | Campeche | 1 | 1 | 0 | 1 | 0 | Edil |
| PB0918 | mxB18 | Campeche | 1 | 1 | 0 | 0 | 1 | Evir |
| PB0921 | mxB18 | Campeche | 1 | 1 | 1 | 1 | 0 | Edil |
| PB0928 | mxB18 | Campeche | 0 | 1 | 0 | 1 | 0 | Edil |
| PB0929 | mxB18 | Campeche | 0 | 1 | 0 | 0 | 1 | Evir |
| PB0933 | mxB18 | Campeche | 0 | 1 | 0 | 1 | 0 | Evir |
| PB0935 | mxB18 | Campeche | 0 | 1 | 0 | 1 | 0 | Edil |
| PB0938 | mxB18 | Campeche | 0 | 1 | 0 | 1R | 0 | Edil |
| PB0940 | mxB18 | Campeche | 0 | 1 | 0 | 1 | 0 | Edil |
| PB0943 | mxB18 | Campeche | 0 | 1 | 0 | 1 | 0 | Edil |
| PB0946 | mxB18 | Campeche | 0 | 1 | 0 | 1 | 0 | Edil |
| PB0947 | mxB18 | Campeche | 0 | 1 | 0 | 1 | 0 | Edil |
| PB0948 | mxB18 | Campeche | 0 | 1 | 0 | 1R | 0 | Evir |
| PB0949 | mxB20 | Morelos | 1 | 1 | 0 | 0 | 1 | Evir |
| PB0950 | mxB20 | Morelos | 1 | 1 | 0 | 0 | 1 | Evir |
| PB0951 | mxB20 | Morelos | 1 | 1 | 0 | 0 | 1 | Evir |
| PB0952 | mxB20 | Morelos | 1 | 1 | 1 | 0 | 1 | Evir |
| PB0953 | mxB20 | Morelos | 1 | 1 | 0 | 0 | 1 | Evir |
| PB0954 | mxB20 | Morelos | 1 | 1 | 0 | 0 | 1 | Evir |
| PB0955 | mxB20 | Morelos | 1 | 1 | 0 | 0 | 1 | Evir |
| PB0956 | mxB20 | Morelos | 1 | 1 | 0 | 0 | 1 | Evir |
| PB0957 | mxB20 | Morelos | 1 | 1 | 0 | 0 | 1 | Evir |
| PB0958 | mxB20 | Morelos | 1 | 1 | 0 | 0 | 1 | Evir |
| PB0959 | mxB20 | Morelos | 1 | 1 | 0 | 0 | 1 | Evir |
| PB0960 | mxB20 | Morelos | 1 | 1 | 0 | 0 | 1 | Evir |
| PB0961 | mxB20 | Morelos | 0 | 1 | 0 | 0 | 1 | Evir |
| PB0962 | mxB20 | Morelos | 0 | 1 | 0 | 0 | 1 | Evir |
| PB0963 | mxB20 | Morelos | 0 | 1 | 0 | 0 | 1 | Evir |
| PB0964 | mxB20 | Morelos | 0 | 1 | 0 | 0 | 1 | Evir |
| PB0965 | mxB20 | Morelos | 0 | 1 | 0 | 0 | 1 | Evir |
| PB0966 | mxB20 | Morelos | 0 | 1 | 0 | 0 | 1 | Evir |
| PB0971 | mxB20 | Morelos | 0 | 1 | 0 | 0 | 1 | Evir |
| PB0976 | mxB20 | Morelos | 0 | 1 | 0 | 0 | 1 | Evir |
| PB0978 | mxB20 | Morelos | 0 | 1 | 0 | 0 | 1 | Evir |
| PB0980 | mxB20 | Morelos | 0 | 1 | 0 | 0 | 1 | Evir |
| SR3363 | mxB13 | Oaxaca | 1 | 0 | 0 | 0 | 1 | Evir |
| SR3369 | mxB13 | Oaxaca | 1 | 1 | 0 | 0 | 1 | Evir |
| SR3371 | mxB13 | Oaxaca | 1 | 1 | 1 | 0 | 1 | Evir |
| SR3372 | mxB13 | Oaxaca | 0 | 1 | 0 | 0 | 1 | Evir |
| SR3374 | mxB13 | Oaxaca | 1 | 1 | 0 | 0 | 1 | Evir |
| SR3375 | mxB13 | Oaxaca | 0 | 1 | 0 | 0 | 1 | Evir |
| SR3378 | mxB13 | Oaxaca | 1 | 1 | 0 | 0 | 0 | Evir |
| SR3379 | mxB13 | Oaxaca | 1 | 1 | 0 | 0 | 1 | Evir |
| SR3380 | mxB13 | Oaxaca | 1 | 1 | 0 | 0 | 1 | Evir |
| SR3382 | mxB13 | Oaxaca | 1 | 1 | 0 | 0 | 1 | Evir |
| SR3383 | mxB13 | Oaxaca | 0 | 1 | 0 | 0 | 1 | Evir |
| SR3384 | mxB13 | Oaxaca | 1 | 1 | 0 | 0 | 1 | Evir |

| | | | | | | | | |
|---|---|---|---|---|---|---|---|---|
| SR3387 | mxB13 | Oaxaca | 1 | 1 | 0 | 0 | 1 | Evir |
| SR3388 | mxB13 | Oaxaca | 0 | 1 | 0 | 0 | 1 | Evir |
| SR3390 | mxB13 | Oaxaca | 1 | 1 | 1 | 0 | 1 | Evir |
| SR3391 | mxB13 | Oaxaca | 0 | 1 | 0 | 0 | 1 | Evir |
| SR3392 | mxB13 | Oaxaca | 0 | 1 | 0 | 0 | 0 | Evir |
| SR3393 | mxB13 | Oaxaca | 0 | 1 | 0 | 0 | 1 | Evir |
| SR3395 | mxB13 | Oaxaca | 0 | 1 | 0 | 0 | 0 | Evir |
| SR3397 | mxB13 | Oaxaca | 0 | 1 | 0 | 0 | 1 | Evir |
| SR3399 | mxB13 | Oaxaca | 0 | 1 | 0 | 0 | 1 | Evir |

**Table S3 Compounds of highest prevalence in the dataset.** Abundance of the 40 most prevalent compounds in the dataset as well as in subsets of *E. dilemma* (Edil) and *E. viridissima* (Evir) are indicated. The last column indicates compounds that contributed more than 1% to perfume differences between species based on a SIMPER analysis (Table S5). For compounds that could not be identified the retention time is indicated.

| Compound | Individuals total | Prevalence total [%] | Individuals Edil | Prevalence Edil [%] | Individuals Evir | Prevalence Evir [%] | SIMPER >1% |
|---|---|---|---|---|---|---|---|
| cis-β-ocimene | 268 | 87.58 | 156 | 84.78 | 112 | 91.80 | x |
| α-pinene | 252 | 82.35 | 142 | 77.17 | 110 | 90.16 | x |
| sabinene | 247 | 80.72 | 143 | 77.72 | 104 | 85.25 | x |
| β-pinene | 234 | 76.47 | 136 | 73.91 | 98 | 80.33 | |
| isoelemicin | 206 | 67.32 | 142 | 77.17 | 64 | 52.46 | x |
| β-caryophyllene | 203 | 66.34 | 124 | 67.39 | 79 | 64.75 | x |
| (-)-terpinen-4-ol | 197 | 64.38 | 112 | 60.87 | 85 | 69.67 | x |
| HNDB4 | 184 | 60.13 | 181 | 98.37 | 3 | 2.46 | x |
| α-caryophyllene | 178 | 58.17 | 109 | 59.24 | 69 | 56.56 | |
| HNDB3 | 176 | 57.52 | 176 | 95.65 | 0 | 0 | x |
| HNDB1 | 171 | 55.88 | 171 | 92.93 | 0 | 0 | x |
| eugenol | 161 | 52.61 | 105 | 57.07 | 56 | 45.90 | x |
| HNDB2 | 159 | 51.96 | 159 | 86.41 | 0 | 0 | |
| cis-β-terpineol | 151 | 49.35 | 90 | 48.91 | 61 | 50.00 | |
| trans-β-ocimene | 148 | 48.37 | 89 | 48.37 | 59 | 48.36 | |
| germacreneD | 146 | 47.71 | 84 | 45.65 | 62 | 50.82 | x |
| linalool | 144 | 47.06 | 93 | 50.54 | 51 | 41.80 | |
| similar to elemicin | 143 | 46.73 | 103 | 55.98 | 40 | 32.79 | |
| τ-terpinen | 141 | 46.08 | 88 | 47.83 | 53 | 43.44 | |
| terpinolen | 138 | 45.10 | 73 | 39.67 | 65 | 53.28 | |
| 1,8-cineol | 137 | 44.77 | 77 | 41.85 | 60 | 49.18 | |
| RT53.4 | 135 | 44.12 | 77 | 41.85 | 58 | 47.54 | |
| D-limonene | 128 | 41.83 | 77 | 41.85 | 51 | 41.80 | x |
| 1-Isopropyl-4-methylbicyclo[3.1.0]hex-2-ene | 125 | 40.85 | 80 | 43.48 | 45 | 36.89 | |
| α-terpinene | 118 | 38.56 | 72 | 39.13 | 46 | 37.70 | |
| α-copaene | 117 | 38.24 | 73 | 39.67 | 44 | 36.07 | |
| tau-elemene | 115 | 37.58 | 66 | 35.87 | 49 | 40.16 | |
| similar to β-bourbonene | 112 | 36.60 | 65 | 35.33 | 47 | 38.52 | |
| similar to β-cubebene | 111 | 36.27 | 72 | 39.13 | 39 | 31.97 | |
| benzyl benzoate | 106 | 34.64 | 93 | 50.54 | 13 | 10.66 | x |
| RT28.7 | 101 | 33.01 | 66 | 35.87 | 35 | 28.69 | x |
| o-cymene | 96 | 31.37 | 49 | 26.63 | 47 | 38.52 | |
| RT26.6 | 96 | 31.37 | 52 | 28.26 | 44 | 36.07 | |
| similar to τ-elemene | 96 | 31.37 | 57 | 30.98 | 39 | 31.97 | |
| similar to thujopsene | 95 | 31.05 | 58 | 31.52 | 37 | 30.33 | |
| L97 | 80 | 26.14 | 5 | 2.717 | 75 | 61.48 | x |
| β-elemene | 77 | 25.16 | 51 | 27.72 | 26 | 21.31 | |
| benzylcinnamate | 76 | 24.84 | 67 | 36.41 | 9 | 7.38 | x |
| 1-terpinen-4-ol | 74 | 24.18 | 50 | 27.17 | 24 | 19.67 | |
| RT27.6 | 72 | 23.53 | 28 | 15.22 | 44 | 36.07 | |

HNDB: 2-hydroxy-6-nona-1,3-dienyl-benzaldehyde; L97: fatty acid lactone derivative of linolenic acid; RT: retention time.

**Table S4 Mean relative abundance of compounds in relation to overall perfume composition.** All compounds with more than 1% relative abundance in either species are presented for *E. dilemma* and *E. viridissima*. The last column indicates compounds that contributed more than 1% to perfume differences between species based on a SIMPER analysis (Table S5). Orange shaded compounds are the major compounds including three stereoisomers of HNDB.

| **Compound** | ***E. dilemma*** | ***E. viridissima*** | **SIMPER>1%** |
|---|---|---|---|
| L97 | 0.04 | 37.28 | x |
| HNDB4 | 48.67 | 0.02 | x |
| HNDB3 | 2.74 | 0.00 | x |
| HNDB1 | 2.73 | 0.00 | x |
| cis-β-ocimene | 5.94 | 8.81 | x |
| benzyl benzoate | 5.90 | 0.07 | x |
| eugenol | 5.17 | 8.27 | x |
| benzylcinnamate | 2.96 | 0.41 | x |
| β-caryophyllene | 2.62 | 3.24 | x |
| isoelemicin | 2.32 | 3.35 | x |
| sabinene | 1.36 | 2.24 | x |
| germacreneD | 1.27 | 3.66 | x |
| D-limonene | 0.92 | 1.95 | x |
| similar to β-cubebene | 0.78 | 1.05 | |
| similar to elemicin | 0.75 | 1.06 | |
| (-)-terpinen-4-ol | 0.69 | 1.97 | x |
| α-pinene | 0.60 | 3.05 | x |
| RT28.7 | 0.53 | 2.79 | x |
| 1,8-cineol | 0.35 | 1.13 | |

HNDB: 2-hydroxy-6-nona-1,3-dienyl-benzaldehyde; L97: fatty acid lactone derivative of linolenic acid; RT: retention time.

**Table S5 SIMPER analysis.** Compounds are ranked by their contribution to chemical dissimilarity of perfumes between *E. dilemma* and *E. viridissima*. Orange shaded compounds are the major compounds including three stereoisomers of HNDB.

| **Compound** | **average contribution to species differences** | **sd** | **Cumulative sum** |
|---|---|---|---|
| HNDB4 | 0.243252592 | 0.110495873 | 0.2622603 |
| L97 | 0.186377211 | 0.200634962 | 0.4632009 |
| eugenol | 0.055175308 | 0.088609215 | 0.5226876 |
| cis-β-ocimene | 0.053654233 | 0.069318033 | 0.5805343 |
| benzyl benzoate | 0.029565737 | 0.048428551 | 0.6124103 |
| isoelemicin | 0.023095267 | 0.049634033 | 0.6373103 |
| β-caryophyllene | 0.021834442 | 0.033611296 | 0.6608508 |
| germacreneD | 0.021382415 | 0.042888817 | 0.6839041 |
| benzylcinnamate | 0.016454031 | 0.040571676 | 0.7016438 |
| RT28.7 | 0.015511842 | 0.03914692 | 0.7183677 |
| α-pinene | 0.015407743 | 0.027173899 | 0.7349794 |
| sabinene | 0.013851618 | 0.025709943 | 0.7499134 |
| HNDB3 | 0.013712336 | 0.009376406 | 0.7646972 |
| HNDB1 | 0.013629247 | 0.011643321 | 0.7793914 |
| D-limonene | 0.012982779 | 0.03795805 | 0.7933887 |
| (-)-terpinen-4-ol | 0.011348614 | 0.040367243 | 0.8056241 |

HNDB: 2-hydroxy-6-nona-1,3-dienyl-benzaldehyde; L97: fatty acid lactone derivative of linolenic acid; RT: retention time.

**Table S6 $f_4$-test for treeness between sympatric and allopatric individuals of *E. dilemma* and *E. viridissima*.**

| Test | $f_4$ | Std. error | Z-score | p-value |
|---|---|---|---|---|
| $f_4(Ed_{allo},Ed_{sym};Ev_{allo},Ev_{sym})$ | 0.00128186 | 0.00047396 | 2.70458 | **0.006841** |

**Table S7 Pairwise $F_{ST}$ between the three genetic lineages $Ed_{south}$, $Ed_{north}$, and *Ev*.** $F_{ST}$ values are shown in the upper right and the number of SNPs used for $F_{ST}$ estimates on the lower left.

| | $Ed_{south}$ | *Ev* | $Ed_{north}$ |
|---|---|---|---|
| $Ed_{south}$ | | 0.04 | 0.09 |
| *Ev* | 4135 | | 0.10 |
| $Ed_{north}$ | 4128 | 4134 | |

**Table S8 Location and size of detected $\Delta F_{ST}$ outlier peaks.** Location of the start and end of each outlier peak on the respective genomic scaffold (*sensu* Brand et al. 2017) are shown.

| Scaffold | Start | End |
|---|---|---|
| scaffold_1 | 150,001 | 2,500,000 |
| scaffold_5 | 4,200,001 | 5,250,000 |
| scaffold_6 | 3,650,001 | 3,700,000 |
| scaffold_28 | 1,700,001 | 1,850,000 |
| scaffold_32 | 50,001 | 1,650,000 |
| scaffold_45 | 1,050,001 | 1,100,000 |
| scaffold_61 | 400,001 | 550,000 |
| scaffold_81 | 150,001 | 200,000 |

**Table S9 Primer for tiled OR41 sequencing.** UTR: Untranslated region.

| Primer | Sequence 5' -> 3' | Product size | Note |
|---|---|---|---|
| OR41.01fwd | CGCCATGTTTCACAAGAGAATG | 846 | starts in 5' UTR; ends in exon 2 |
| OR41.01rev | GGTACAGGTTGTTGCACGAG | | |
| OR41.02fwd | TCTCACGCGTATTTCTATG | 393 | starts in intron 1; ends in exon 3 |
| OR41.02rev | AGTTACATTCCTTTCCCTGTTTA | | |
| OR41.03fwd | GAGATTATGCACCACCCTC | 836 | starts in exon 2; ends in intron 4 |
| OR41.03rev | ACGGTATTACTTAAAACGTATCG | | |
| OR41.04fwd | GAATTATTGCAAGAATCTGGAG | 885 | starts in exon 4; ends in 3' UTR |
| OR41.04rev | ACCGTCATTGTGAGAAATATCA | | |

**Table S10 Individuals sequenced for *OR41*.**

| Sample | Location | Genotype |
|---|---|---|
| PB0216 | crB04 | $Ed_{south}$ |
| PB0473 | mxB05 | $Ed_{south}$ |
| PB0496 | mxB05 | $Ed_{south}$ |
| PB0526 | mxB06 | $Ed_{south}$ |
| PB0541 | mxB06 | $Ed_{south}$ |
| PB0545 | mxB06 | $Ed_{south}$ |
| PB0561 | mxB08 | *Ev* |
| PB0564 | mxB08 | *Ev* |

| | | |
|---|---|---|
| PB0566 | mxB08 | $Ed_{north}$ |
| PB0589 | mxB08 | $Ed_{north}$ |
| PB0596 | mxB09 | *Ev* |
| PB0601 | mxB09 | $Ed_{north}$ |
| PB0618 | mxB09 | $Ed_{north}$ |
| PB0682 | mxB12 | *Ev* |
| PB0685 | mxB12 | *Ev* |
| PB0698 | ho01 | $Ed_{south}$ |
| PB0699 | ho01 | $Ed_{south}$ |
| PB0700 | ho01 | $Ed_{south}$ |
| PB0705 | ho01 | $Ed_{south}$ |
| PB0729 | mxB15 | *Ev* |
| PB0758 | mxB15 | *Ev* |
| PB0760 | mxB15 | *Ev* |
| PB0761 | mxB15 | *Ev* |
| PB0763 | mxB15 | $Ed_{north}$ |
| PB0769 | mxB15 | $Ed_{north}$ |
| PB0786 | mxB16 | $Ed_{north}$ |
| PB0810 | mxB17 | $Ed_{north}$ |
| PB0844 | mxB17 | $Ed_{north}$ |
| SR640 | gtB01 | $Ed_{north}$ |
| SR656 | gtB01 | $Ed_{south}$ |
| SR2292 | Costa Rica | $Ed_{south}$ |
| SR2295 | Costa Rica | $Ed_{south}$ |
| SR2303 | Florida | $Ed_{south}$ |
| SR2342 | Nicaragua | $Ed_{south}$ |
| SR2355 | Nicaragua | $Ed_{south}$ |
| SR2365 | Nicaragua | $Ed_{south}$ |
| SR2432 | Florida | $Ed_{south}$ |
| CD136 | usB01 | $Ed_{south}$ |
| CD139 | usB01 | $Ed_{south}$ |
| CD156 | Florida | $Ed_{south}$ |
| CD160 | Costa Rica | $Ed_{south}$ |
| CD177 | Mexico | $Ed_{north}$ |
| CD183 | Mexico | *Ev* |
| CD213 | Mexico | *Ev* |
| CD243 | Mexico | $Ed_{north}$ |
| CD250 | Mexico | *Ev* |
| CD269 | Mexico | $Ed_{north}$ |

**Table S11 Selection test based on the *OR41* phylogeny.** The *OR41* phylogeny was tested with *E. dilemma* as foreground and *E. viridissima*, *E. imperialis*, *E. flammea*, and *Ef. mexicana* as background.

| Model | $d_N/d_S$ background | $d_N/d_S$ foreground | log-likelihood |
|---|---|---|---|
| background = foreground | 0.3969 | 0.3969 | -2482.280816 |
| background ≠ foreground | 0.321 | 3.6283 | -2474.225479 |
